# A Micro-Engineered Heart Tissue Model of Desmin-related Cardiomyopathy Caused by Mutant αB Crystallin

**DOI:** 10.1101/2025.10.02.680018

**Authors:** Yasaman Kargar Gaz Kooh, Bahareh Bahmani, Chen Zhao, Ganesh Malayath, Leah Hayem, Hanxun Jin, Ghiska Ramahdita, Huanzhu Jiang, Javier Santiago Perez, Hsin Yi Chou, Xiucui Ma, Guy Genin, David Rawnsley, Abhinav Diwan, Nathaniel Huebsch

**Author notes:** **Corresponding Author:** Nathaniel Huebsch, PhD. Whitaker Hall 300E 1 Brookings Drive Saint Louis, MO, 63130.

## Abstract

Protein quality control (PQC) is essential for maintaining sarcomere integrity in cardiomyocytes. α Crystallin B chain (CRYAB) R120G mutation disrupts CRYAB’s chaperone activity, leading to aggregation of CRYAB and its client proteins (including Desmin), leading to Desmin-related cardiomyopathy (DRM). Prior experimental systems for modeling DRM linked to CRYAB require massive overexpression of CRYAB mutant isoforms, raising questions about translational relevance. Here, we establish the first model of CRYAB-linked DRM that uses genome-edited hiPSC together with isogenic controls, allowing us to study the impact of mutant CRYAB expressed at near endogenous levels. Within micro-engineered heart tissues (μHT), CRYAB-R120G mutant hiPSC-derived cardiomyocytes recapitulated key DRM hallmarks, including Desmin and CRYAB aggregation, contractile dysfunction, and increased vulnerability to PQC pathway inhibition. CRYAB-R120G mutant μHT also exhibited dysfunctional calcium-contraction coupling, which exacerbated contractile deficits at higher pacing frequencies. JAK1 inhibition with Itacitinib partially restored contractile function at higher pacing frequencies, suggesting JAK1 inhibition as a viable therapeutic strategy. By preserving human-specific structural and functional features, our µHT platform enables mechanistic characterization of proteotoxic cardiomyopathies and offers a scalable system for targeted drug screening.

## Introduction

Heart failure remains a leading cause of death worldwide (1). Inherited cardiomyopathies contribute significantly to early onset heart failure (2). Desmin-related cardiomyopathy (DRM) is a form of familial cardiomyopathy that affects approximately 1 in 2000 individuals (3). It is caused by mutations in key cytoskeletal components such as Desmin (4, 5) or its chaperone α-Crystallin B chain (CRYAB) (6, 7). Among DRM-linked CRYAB mutations, the missense R120G variant, which is inherited in an autosomal dominant fashion, is especially pathogenic. This mutation promotes toxic aggregation of both CRYAB and Desmin, leading to sarcomere disorganization, proteotoxic stress, and impaired contractile function (6–8).

DRM pathology is a striking example of dysfunctional cardiomyocyte protein quality control (PQC). As “sarcomeres are machines that must repair themselves while in motion (9)”, cardiomyocytes depend on highly regulated PQC networks to degrade or repair proteins that become misfolded through genetic mutations or mechanical/oxidative stress (9, 10). Chaperones like CRYAB first attempt to refold these misfolded proteins. Those that cannot be refolded are tagged with poly-ubiquitin chains and sent to the ubiquitin-proteasome system for degradation. The autophagy-lysosome pathway provides a second backup, where ubiquitinated proteins and damaged organelles are taken up by autophagosomes that fuse with lysosomes for final clearance. Together, these complementary PQC pathways preserve cardiomyocyte proteostasis (9, 11–14).

Failure of PQC, linked to genetic mutations or chemical stress induced by drugs like proteosome inhibitors used for chemotherapy, causes cardiomyopathy (15, 16). This cardiomyopathy occurs because accumulation of cytosolic aggregates of misfolded proteins overwhelms the cellular PQC machinery, preventing *in situ* repair of the sarcomeres; this leads to contractile failure and eventually cell death (17, 18). Proteotoxicity of Desmin aggregates is a hallmark of DRM (18). Desmin, a muscle-specific intermediate filament protein that anchors sarcomeres to organelles such as mitochondria to maintain structural integrity and force transmission, is especially dependent on PQC (19, 20). Beyond DRM linked to mutations and chemotherapy, Desmin aggregation has been linked to cardiac ischemia (21), highlighting the potential broad relevance of understanding the pathogenesis of DRM and developing treatments.

DRM is strongly linked to the dysfunction of CRYAB, a small heat shock protein that is abundantly expressed in cardiac muscle which acts as a molecular chaperone for sarcomere proteins including Desmin, preventing misfolding and guiding these proteins to their final location in the sarcomere (Desmin itself resides in the Z-disc (21, 22)). The R120G mutation in CRYAB is especially toxic because it weakens both the protein’s structural stability and its chaperone activity. Computational (23) and experimental studies (24, 25) show that the R120G mutation breaks key stabilizing salt bridges (R120-D109 and R107-D80), collapsing the central cavity of the α-crystallin domain, to expose hydrophobic regions by disrupting the IPI/β4-β8 interaction. This leaves unpaired negative charge that introduces an overall electrostatic imbalance. This altered surface charge likely makes mutant CRYAB bind more tightly to Desmin, trapping Desmin into a misfolded complex (18), and driving formation of large and insoluble oligomers consisting of both Desmin and CRYAB. These aggregates, in turn, can sequester other client proteins, leading to marked cellular dysfunction.

Despite the clinical relevance of genetically inherited DRM and closely related drug and ischemia-induced DRM, mechanisms underlying DRM remain poorly understood, and medical management remains limited to symptomatic therapies such as heart failure medications (e.g., β-blockers, ACE inhibitors) and antiarrhythmics, as there are no disease-specific therapies targeting the protein aggregation pathology (7). Previous studies investigating CRYAB mutations like R120G have relied on mouse models with transgene-mediated overexpression of the mutant isoforms of this protein (18, 26–28). This raises two important concerns: First, as CRYAB itself tends to be found aggregated in DRM, and the requirement for massive overexpression (typically 6-fold increase in absolute CRYAB level is required to induce cardiomyopathy phenotypes in mice (27)) raises concerns about physiological relevance given the potential for overexpression to induce aggregation as an experimental artifact. Second, in addition to concerns involving requirements for high absolute CRYAB expression levels, the reliance on murine systems raises questions about translatability to human cardiomyocytes (29), particularly given reported differences in basal and stress-induced expression of heat shock proteins, such as HSP70, between humans and rodents (30).

Cell culture-based model systems using human iPSC-derived cardiomyocytes (hiPS-CMs) offer a species-relevant platform for studying cardiomyopathy (29). However, hiPS-CM are often structurally and functionally immature, which has limited their application for disease modeling (31–35). To overcome these limitations, we developed a 3D micro-engineered heart tissue (µHT) platform using hiPS-CMs derived either from hiPSC engineered to harbor a homozygous knock-in of the CRYAB-R120G mutation, or isogenic controls. This allowed us to study pathology associated with CRYAB mutations at near-physiologic expression levels. To our knowledge, this is the first hiPSC model of this mutation with near physiologic protein expression and paired with an isogenic control, allowing mechanistic studies.

Because Desmin expression was poor in our hiPS-CM in baseline, we first compared maturation-inducing chemical treatments to culture within 3D engineered µHT as means of inducing physiologic Desmin expression and localization (36, 37). Strikingly, only hiPS-CM within the µHT exhibited robust expression, with appropriate physiological localization to the Z-disc, of this key intermediate filament protein. As in prior murine models of DRM linked to CRYAB-R120 overexpression (18, 28), the knock-in R120G mutation in CRYAB caused marked contractile dysfunction in μHT. Studies with voltage dyes and the genetically encoded calcium indicator RGECO1.2 revealed that this contractile dysfunction was linked to dysregulated calcium-contraction coupling, which made contractility deficits especially pronounced at higher pacing frequencies. Functional pathology associated with the CRYAB-R120G mutation was associated with aggregation in CRYAB and its client, Desmin. This proteotoxic stress has been shown to activate inflammatory signaling cascades, especially JAK/STAT (38). Thus, we evaluated the JAK1-specific inhibitor, Itacitinib, as a candidate for DRM therapy. This treatment partially rescued the contractility deficits associated with the CRYAB-R120G mutant μHT. These findings establish the first cell culture-based model of DRM with physiologically relevant CRYAB-R120G expression, enabling further mechanistic investigation and drug screening of DRM in a human context.

## Results

### Development of a high-throughput hiPS-CM based platform to study pathology associated with the CRYAB-R120G point mutation

In postnatal cardiomyocytes, Desmin is abundantly expressed and is primarily localized to desmosomes and Z-discs (39, 40). At baseline, we observed vanishingly low levels of Desmin expression in hiPS-CM, with only about 1 in 100 cells expressing this protein. This is consistent with prior reports that deficient expression and mislocalization of Desmin make DRM especially challenging to model in immature hiPS-CMs (35). We thus evaluated two strategies for enhancing Desmin expression in hiPS-CM: (1) culturing cells in monolayer using a defined metabolic maturation media (MM) (41, 42), and (2) co-culturing cells with primary cardiac fibroblasts and extracellular matrix (ECM) under mechanical stimulation within 3D µHT (**Supplemental Video 1**).

We first assessed a MM that uses fatty acids and the PPAR-δ agonist (GW0742) to promote hiPS-CM structural maturation in 2D culture (42) (**Fig. S1 A, B**). Analysis of cardiomyocyte morphology facilitated by Cellpose (43) (**Fig. S1 C**), an unbiased and AI-assisted segmentation package, confirmed that MM-induced structural maturation (significant cellular elongation with no significant cellular hypertrophy) when compared against standard media (SM; RPMI-1640 supplemented with 2% B27 supplement) (**Fig. S1 D-G**). This is consistent with structural maturation previously observed for monolayers cultured in this same MM (42). However, even with this structural maturation, the overall number of hiPS-CM expressing Desmin was still vanishingly small (∼1%) (**see below**).

Given the lack of Desmin expression in monolayer cultures, we transitioned to a tissue engineering approach (44). We formed μHT by co-encapsulating 95% hiPS-CMs with 5% primary human cardiac fibroblasts into a mixture of exogenous ECM proteins within micro-wells containing PDMS pillars (**Fig. 1 A, B; Fig. S2**).

**Figure 1.**
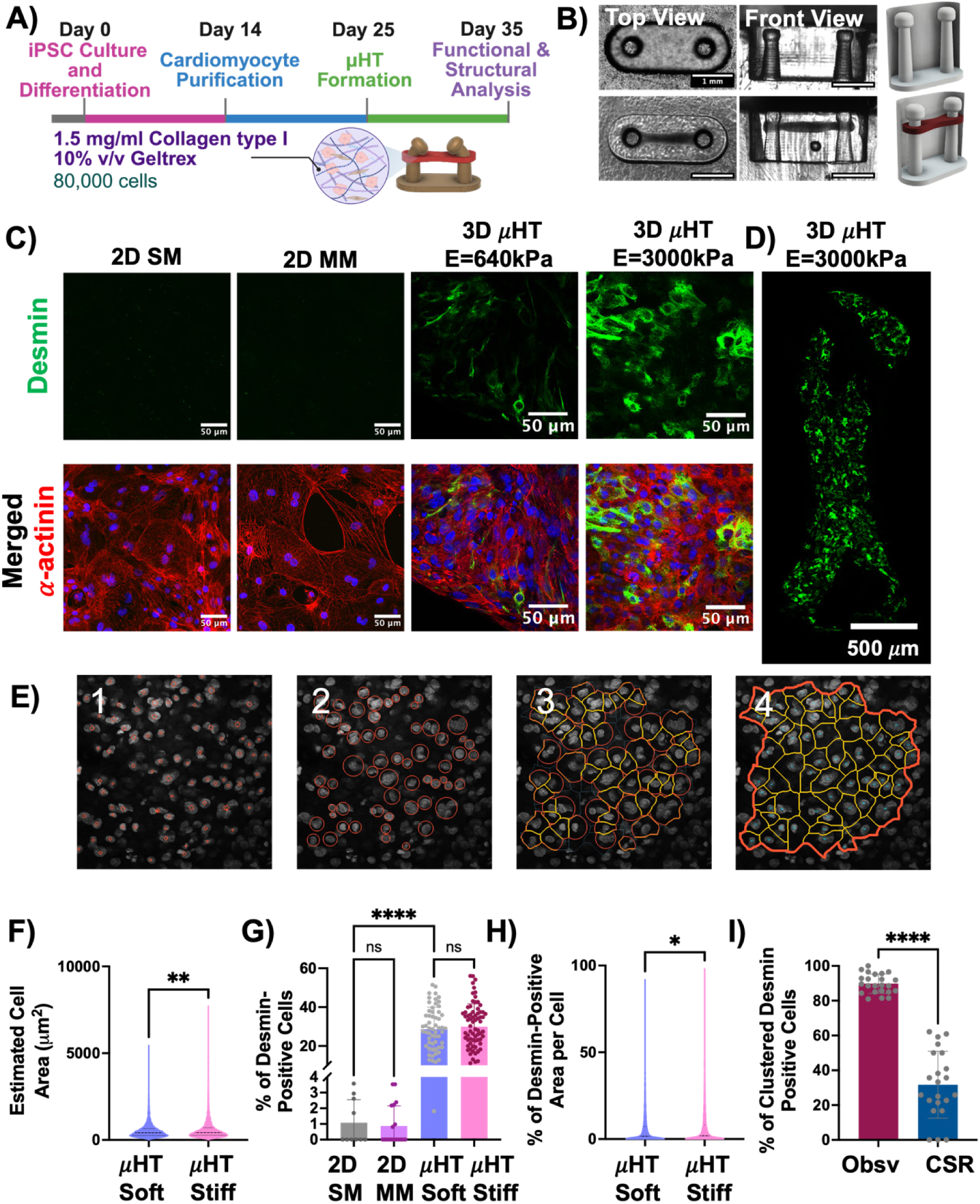
Culture within 3D μHT triggers Desmin expression in hiPS-CM. **(A)** Schematic of the experimental timeline for generating 3D μHT from hiPS-CMs. Cells were differentiated from hiPSCs (D0–D14), purified by metabolic selection (D14–D25), and seeded into 3D μHT devices for culture from D25–D35 before analysis. **(B)** Representative brightfield images of μHTs from top and front views on PDMS posts, alongside schematic micrograph of the post geometry with and without tissue load. Scale bars: 1mm. **(C)** Immunofluorescence staining for α-actinin (red), Desmin (green), and Hoechst (blue) in either 2D with SM or MM and μHTs cultured on soft (E = 640 kPa) or stiff (E = 3000 kPa) posts. Scale bars: 50 μm. **(D)** Wide-field view of the whole-mount immunostaining for Desmin in 3D μHTs cultured on stiff posts. Scale bar: 500 μm. **(E)** Representative images showing crystal growth from nuclear segmentation for cell shape analysis: raw Hoechst stain (1), nuclei with red outlines (2), enlarging the cell boundary (3), and overlaid segmentation boundaries (4). **(F)** Quantification of estimated cell area from nuclei-stained μHTs, showing significantly increased cell area on stiff posts compared to soft posts. Data are presented as mean ± *SD*; n values represent individual cells pooled from 3 independent differentiation batches; *P* values by unpaired Student t-test, non-parametric, Mann-Whitney test; ** *p* < 0.01. **(G)** Quantification of the percentage of Desmin-positive cells comparing 2D cultured in either SM or MM with 3D μHTs within soft and stiff posts. Data are presented as mean ± *SD*; n values represent individual 2D monolayers or individual μHTs pooled from 3 independent differentiation batches; *P* values by one-way ANOVA, non-parametric, Kruskal Wallis test; **** *p* < 0.0001, ns = not significant. **(H)** Quantification of the percentage of Desmin-positive area per cell in 3D μHTs, revealing a significant increase in Desmin-positive area per cell on stiff posts. Data are presented as mean ± *SD*; n values represent individual cells pooled from 3 independent differentiation batches. *P* values by unpaired t-test, non-parametric, Mann-Whitney test; * *p* < 0.05. **(I)** Quantification of the percentage of clustered Desmin-positive cells in 3D μHTs compared to simulations of complete spatial randomness (CSR). Observed (Obsv) distributions showed significantly higher clustering than CSR, supporting a non-random and localized expression of Desmin-positive hiPS-CMs. Data are presented as mean ± *SD*; n values represent individual μHTs pooled from 3 independent differentiation batches. *P* values by unpaired Student t-test, parametric **** *p* < 0.0001.

Strikingly, even without maturation-inducing chemicals such as fatty acids, within µHTs formed against compliant PDMS posts (E = 640 kPa, bending stiffness of 0.42 N/m corresponding to applied force of 98 µN at 234 µm deflection, as estimated from finite element simulations), hiPS-CM exhibited robust Desmin expression (**Fig. 1 C)**. Consistent with our prior studies on Desmin staining in positive control postnatal human heart sections (21), Desmin in hiPS-CM within μHT was co-localized with α-actinin at Z-discs (**Fig. 1 C**). Interestingly, within µHTs formed against stiff PDMS posts (E = 3000 kPa, estimated bending stiffness: 1.96 N/m corresponding to applied force of 98 µN at 50 µm deflection; **Fig, S3 B**), hiPS-CM showed even more robust Desmin expression, and similar Z-disc localization (**Fig. 1 C**).

Whole-mount immunostaining followed by high-resolution image stitching of µHTs labeled for α-actinin, Desmin and Hoechst allowed visualization of the entire tissue construct and localization patterns of Desmin within a single field of view. In this context, Desmin expression appeared broadly uniform throughout the tissue, with no obvious regional bias between the central shaft and areas close to the PDMS posts (**Fig. 1 D; Fig. S4)**. These results suggest that Desmin expression is independent of local stress amplitude (45, 46) or stress anisotropy (36, 44).

To gain insights into whether culture in μHT caused a uniform, per-cardiomyocyte increase in Desmin, or if Desmin expression was instead driven by an “elite” sub-population of cells, we again turned to AI-assisted cell segmentation. Unambiguous, automated identification of cell boundaries in the stained μHT, even with membrane-specific stains like wheat germ agglutinin (WGA) proved challenging **(Fig. S5)**. Moreover, cortical staining for α-actinin, which was highly prominent in 2D monolayers (**Fig. 1 C; Fig. S1 B)**, was not strongly apparent in 3D μHT, making it difficult even for trained observers to identify individual cells (47). We thus developed a custom image segmentation approach designed for dense tissue environments (**Supplement Video 2, Fig. 1 E**). In this method, individual cell boundaries are estimated from nuclear staining alone; the approach exploits the general observation that even in high density 3D tissues like μHT, nuclei can typically be identified in an unbiased and unambiguous manner. Because of the high cell density of μHT (**Fig. 1 C**), we assumed that each cell “grows out” from the nucleus to occupy space until it impinges on the borders of its neighbors. Algorithmically, we accomplished this with an approach inspired by traditional watershed segmentation and the growth of crystals from the melt of crystalline phase materials (48). First, a circular border was applied to each identified nucleus. Next, each border was allowed to expand until they impinged upon the expanding border of its nearest neighbors (**Fig. 1 E)**. The estimated projected area and shape of each cell was then quantified.

Morphological estimations suggested a significant increase in overall hiPS-CM area within µHTs formed against stiff posts (**Fig. 1 F**), consistent with prior observations by our group and others that hiPS-CM undergo hypertrophy when working against a higher afterload (49–51). However, this analysis revealed no correlated changes in estimated eccentricity (**Fig. S6 A**), elongation (**Fig. S6 B**), or circularity (**Fig. S6 C**).

Finally, we measured Desmin expression in each identified cell. Remarkably, this analysis revealed that on average, ∼30% of hiPS-CM in a given μHT region of interest expressed Desmin, regardless of post rigidity (**Fig. 1 G**). While the percentage of Desmin-positive cells was similar in μHT working against posts with different stiffness (**Fig. 1 G**), the percent of Desmin-positive area in each identified cell was significantly higher in µHTs working against a higher afterload (**Fig. 1 H**). This suggests that afterload is a key determinant of hiPS-CM Desmin expression. A similar trend was observed in mutant CRYAB-R120G µHTs, which exhibited increased estimated cell area (**Fig. S7 A**), but reduced elongation (**Fig. S7 B**), eccentricity (**Fig. S7 C**), and no change in circularity (**Fig. S7 D**). As in the isogenic control, the percent of Desmin-positive cells was comparable across conditions (**Fig. S7 E**); however, the percent of Desmin-positive area per cell was significantly greater in µHTs working against stiffer posts (**Fig. S7 F**).

On a small scale, Desmin expression appeared to follow a distinct pattern: hiPS-CM positive for Desmin were mostly found in clusters, close to one another, rather than being randomly distributed throughout the tissue. To demonstrate this clustering pattern in a robust manner, we performed computational simulations to ask how often, in a population where 30% of cells are Desmin-positive, would we expect to find the Desmin-positive cells in larger clusters (>2 cells/cluster) if the cells were distributed randomly (**Fig. S8**). This analysis confirmed our observation that indeed, the Desmin-positive hiPS-CM within μHT tend to group into large cluster (**Fig. 1 I**).

Comparison of observed versus simulated distributions of Desmin-positive cells within the µHT confirmed the non-random, clustered spatial distribution of Desmin-positive cardiomyocytes (**Fig. 1 I; Fig. S8 A-C**). Analysis of 25 randomly selected clusters of Desmin-positive cells indicated that on average, cluster circularity was 0.37 ± 0.11, while eccentricity was 0.85.± 0.1. These quantitative data support our qualitative observation that clusters of Desmin-positive cardiomyocytes showed an elongated, rather than circular shape, suggesting that directional mechanical forces within the μHT may guide the alignment of Desmin-positive cells (**Fig. S8 D**). Altogether, these observations suggest the possibility that localized paracrine signaling and/or localized changes in rigidity of the fibrous tissues are means through which afterload enhances Desmin expression (52).

Overall, these results suggest that while metabolic maturation media can improve some aspects of hiPS-CM structural maturation in 2D culture, it is not sufficient to induce Desmin expression. In contrast, forming 3D µHT with mechanical loading leads to robust Desmin expression.

### hiPS-CMs harboring the CRYAB-R120G point mutation exhibit Desmin aggregation

Having demonstrated an ability to assess Desmin structure within μHT, we next asked whether human cardiac myocytes harboring the DRM-associated R120G mutation in CRYAB would exhibit Desmin abnormalities without a requirement for massive overexpression of the mutant protein. Using CRISPR/Cas9, we generated an isogenic hiPSC line homozygous for the CRYAB-R120G mutation. Within μHT, hiPS-CMs harboring mutant CRYAB showed similar levels of Desmin-positive cells (**Fig. 2 A; Fig. S9 A**) and Desmin-positive area (**Fig. 2 B**) compared to hiPS-CM in isogenic control μHT. Strikingly, CRYAB mutant tissues showed significant structural disorganization and aggregation of Desmin. Quantitatively, this was related to similar number of Desmin-positive cells in aggregated form (**Fig S9 B**) but with a markedly increased area of hiPS-CM expressing Desmin in an aggregated form rather than Z-disc-localized pattern (**Fig. 2 C).** Moreover, consistent with reports that transgenic R120G mice develop cardiac hypertrophy (28), DRM mutant µHTs exhibited a statistically significant increase in estimated cardiomyocyte size (**Fig. 2 D)**.

**Figure 2.**
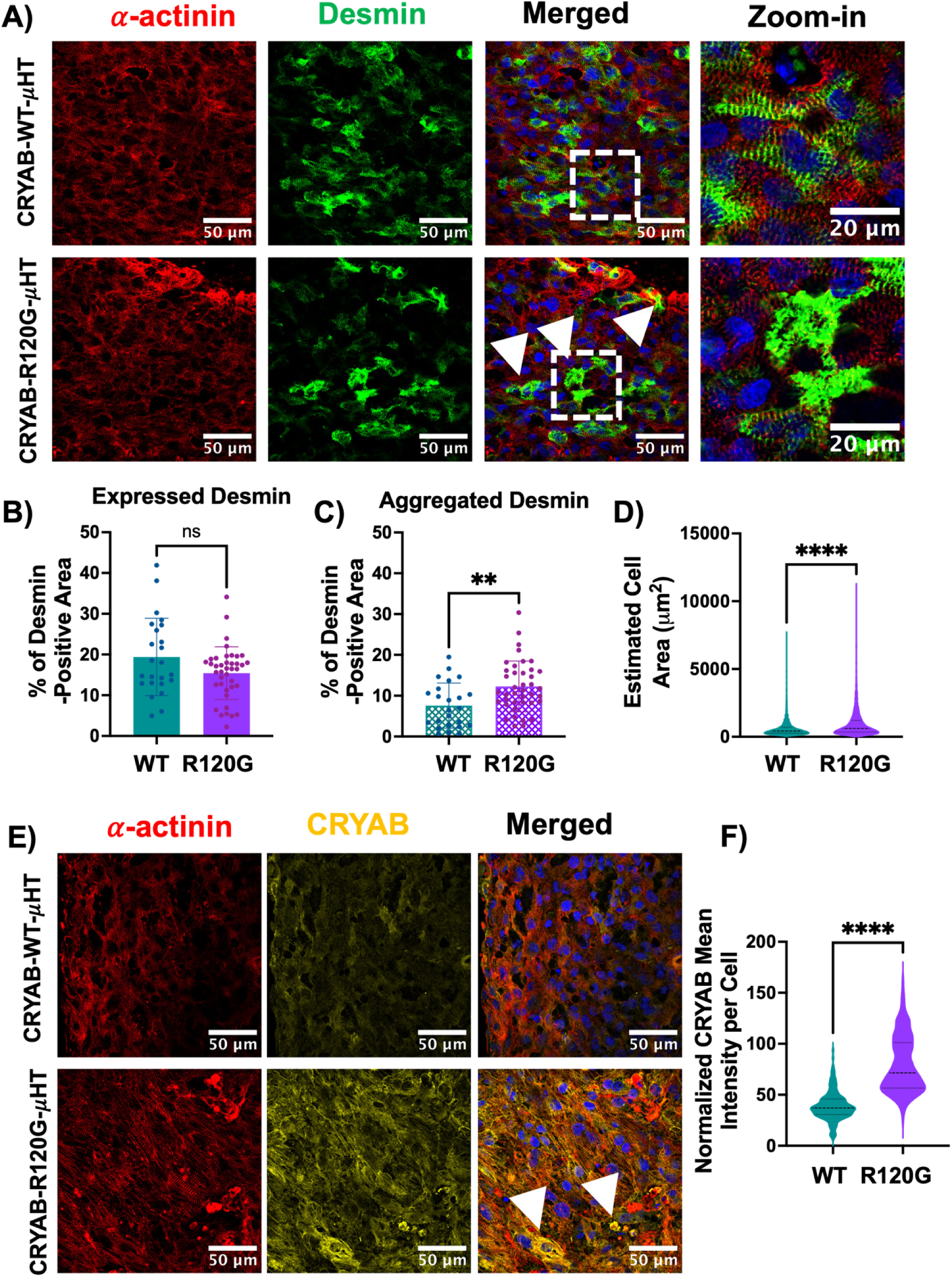
CRYAB-R120G mutation promotes Desmin aggregation and increases CRYAB accumulation in 3D μHT. **(A)** Immunofluorescence staining for α-actinin (red), Desmin (green), and Hoechst (blue) in CRYAB-WT-μHT and CRYAB-R120G-μHT at day 35. Arrows indicate regions of Desmin aggregation in CRYAB-R120G-μHTs. Scale bars: 50 μm (merged images), 20 μm (zoom-in). **(B, C)** Quantification of Desmin expression and aggregation in μHTs. **(B)** The percentage of total tissue area occupied by Desmin-positive staining was measured across μHTs. No significant difference was observed between WT and CRYAB-R120G μHTs. **(C)** The percentage of Desmin-positive area in aggregation form was calculated relative to the total Desmin-positive area. CRYAB-R120G μHTs showed a significantly higher proportion of aggregated Desmin compared to WT. Data are presented as mean ± *SD*; n values represent individual μHT pooled from 3 independent differentiation batches. *P* values were determined by unpaired Student’s t-test, parametric; ** *p* < 0.01. **(D)** Quantification of estimated cell area from nuclei-stained μHTs demonstrate significantly higher estimated cell area in CRYAB-R120G mutant μHTs compared to isogenic controls, consistent with a hypertrophic phenotype in this mutation. Data are presented as mean ± *SD*; n values represent individual cells pooled from 3 independent differentiation batches. *P* values were determined by unpaired Student’s t-test, non-parametric, Mann–Whitney test; **** *p* < 0.0001. **(E)** Immunofluorescence staining for α-actinin (red), CRYAB (yellow), and Hoechst (blue) in CRYAB-WT-μHT and CRYAB-R120G-μHT. Arrow indicates region of CRYAB accumulation in CRYAB-R120G-μHTs. Scale bars: 50 μm. **(F)** Quantification of mean CRYAB fluorescence intensity per cell, showing a significant increase in CRYAB levels in CRYAB-R120G-μHTs compared to isogenic controls. Data are presented as mean ± *SD*; n values represent individual cells pooled from ≥3 independent differentiations. *P* values by unpaired t-test, non-parametric, Mann-Whitney test; **** *p* < 0.0001.

Mutations in CRYAB, including R120G, have been associated with disruptions of this protein’s chaperone functions, thereby leading to aggregation of client proteins like Desmin, but also with aggregation of CRYAB itself (6). We thus examined CRYAB levels and localization within the μHT. Strikingly, we observed that the mean CRYAB expression per cell was elevated nearly 2-fold in hiPS-CM harboring the R120G mutation (**Fig. 2 E, F**). This may result from inhibited clearance of aggregate-associated CRYAB-R120G, leading to an effective increase in steady-state levels of this protein.

### CRYAB-R120G mutant µHTs exhibit contractile deficits

Under low afterload conditions (soft posts; **Fig. 3 A**), CRYAB-R120G µHTs showed significant deficits in contractile performance, as illustrated by representative force traces highlighting disrupted force development and slower contractile kinetics compared to isogenic controls (**Supplemental Video 3**; **Fig. 3 B**). This was reflected in depressed peak active force (**Fig. 3 C**) and reduced relaxation speed (**Fig. 3 D**). Importantly, these deficits persisted under high afterload conditions (stiff posts; **Fig. 3 E**), where CRYAB-R120G (**Supplemental Video 4**) µHTs again showed significantly reduced peak active force and relaxation speed (**Fig. 3 F-H**). Quantitative analysis of additional contraction parameters, summarized in the heatmap (**Fig. 3 I, Fig. S10**), further supports the marked hypocontractile phenotype in CRYAB mutant µHTs. Mutant µHTs showed slower contraction and relaxation speeds, lower active stress, and lower peak deflections compared to isogenic control, regardless of post stiffness.

**Figure 3.**
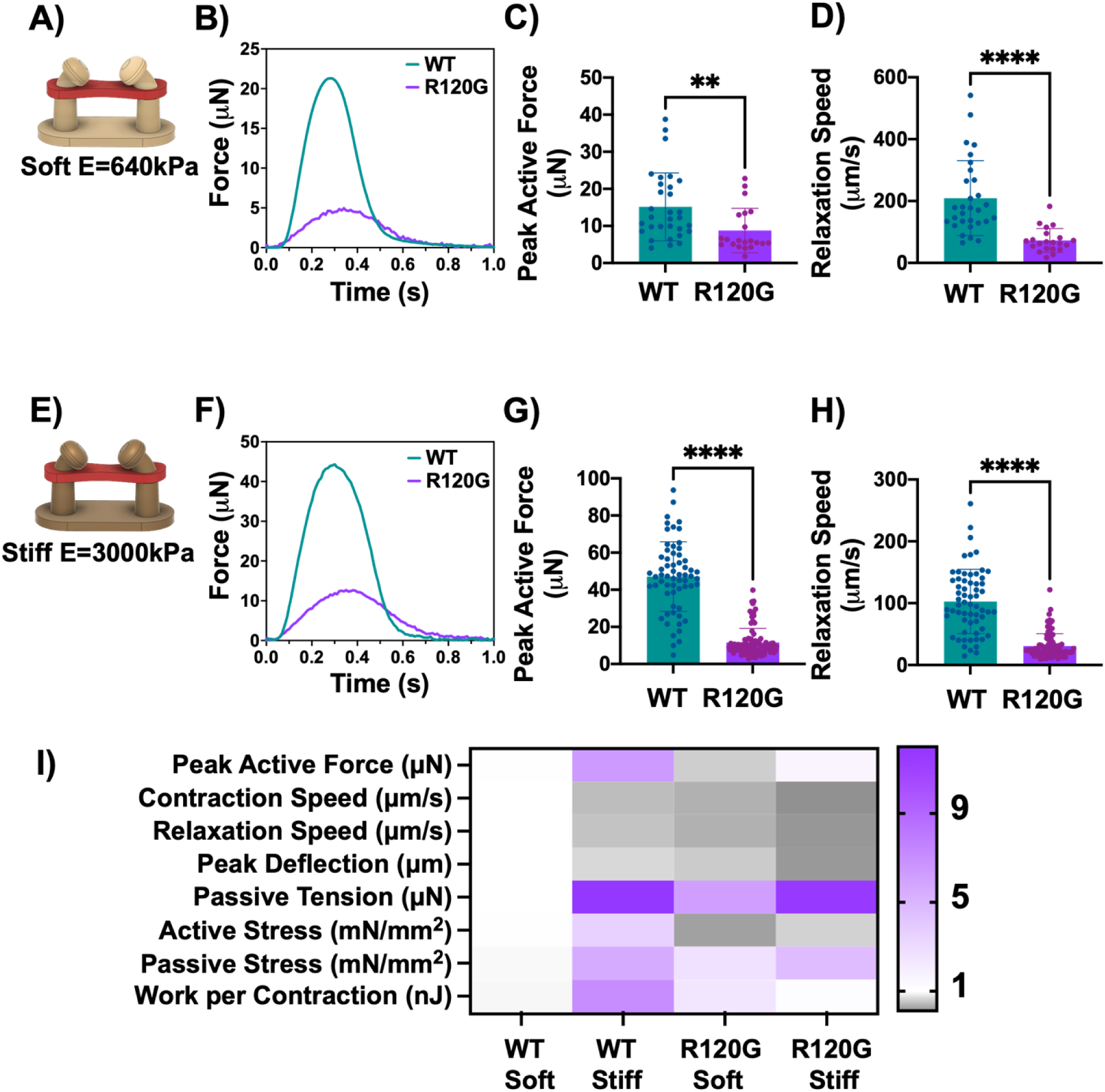
CRYAB-R120G mutation reduces contractile performance of 3D μHTs. (A-D) Soft post condition (E = 640 kPa). **(A)** Schematic of 3D μHT on soft PDMS posts. **(B)** Representative force–time traces for CRYAB-WT and CRYAB-R120G μHTs. Quantification of **(C)** peak active force and **(D)** relaxation speed, showing significantly impaired contractility in CRYAB-R120G μHTs compared to isogenic controls. Data are presented as mean ± *SD*; n values represent individual μHT pooled from ≥3 independent differentiations. *P* values by unpaired t-test, non-parametric, Mann-Whitney test; **** *p* < 0.0001, ** *p* < 0.01. **(E-H)** Stiff post condition (E = 3000 kPa). **(E)** Schematic of 3D μHT on stiff PDMS posts. **(F)** Representative force–time traces for CRYAB-WT and CRYAB-R120G μHTs. Quantification of **(G)** peak active force and **(H)** relaxation speed, showing a pronounced contractility deficit in CRYAB-R120G μHTs similar to μHT cultured within soft posts. Data are presented as mean ± *SD*; n values represent individual μHT pooled from ≥3 independent differentiations. *P* values by unpaired t-test, non-parametric, Mann-Whitney test; **** *p* < 0.0001. **(I)** Heatmap of contractile and mechanical parameters across conditions, including peak active force, contraction speed, relaxation speed, peak deflection, passive tension, active and passive stress, and peak work per contraction cycle. All values were normalized to the mean value of the WT soft condition, with darker purple indicating higher normalized values.

Given that stiffer post conditions promoted greater hiPS-CM Desmin expression consistent with prior findings that afterload enhances maturation of engineered heart tissues (49), along with physiologic hypertrophy, more robust sarcomere organization (**Fig. 1**), and higher peak active force compared to soft posts, we prioritized the high afterload (elastic modulus: 3000 kPa) posts for subsequent analyses.

### CRYAB-R120G mutation make μHT more vulnerable to disruptions in protein quality control pathways

We next tested whether disrupting the heat shock protein system via the CRYAB-R120G mutation would make μHT more vulnerable to inhibition of either of the two alternative PQC pathways, the ubiquitin-proteasome system and lysosome-autophagy. To assess vulnerability to ubiquitin-proteasome system inhibition, we treated µHTs with Bortezomib (100 nM) for 48 hours **(Fig. 4 A)**. Immunofluorescence analysis of Desmin and α-actinin following Bortezomib exposure further supported this vulnerability. While both isogenic control and CRYAB-R120G µHTs exhibited Desmin aggregation after proteasome inhibition **(Fig. 4 B)**, sarcomeric α-actinin striation patterns remained relatively preserved in isogenic controls. In contrast, CRYAB-R120G µHTs displayed extensive sarcomere disorganization, with poorly aligned myofibrils. Contractility was measured in the same tissues before and after treatment, and the contractility index was defined as the ratio of post-treatment to pre-treatment peak active force. Consistent with our expectation, CRYAB-R120G µHTs showed a significantly greater reduction in contractile index following Bortezomib exposure compared to isogenic controls **(Fig. 4 D).**

**Figure 4.**
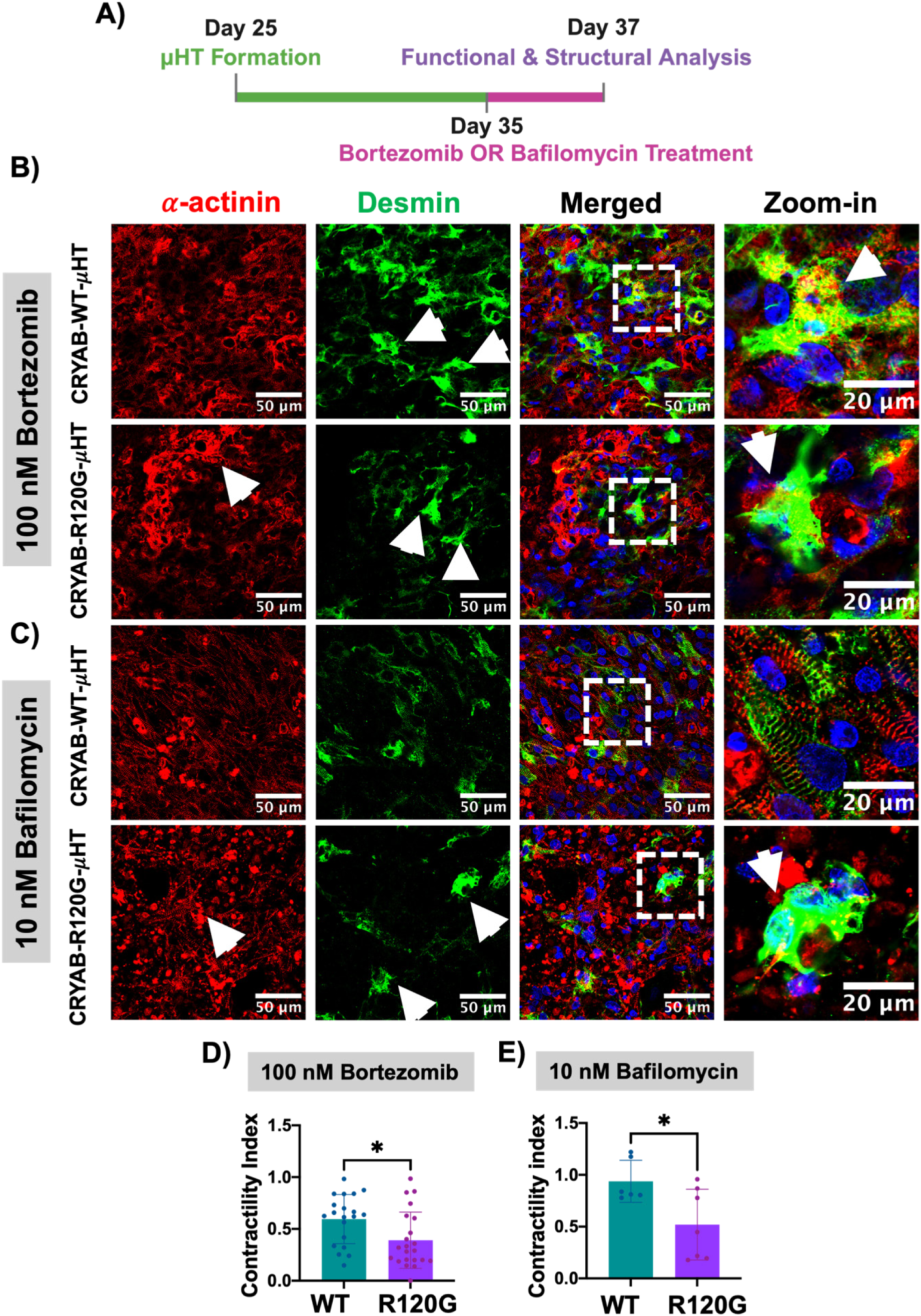
The CRYAB-R120G mutation increases vulnerability of μHT to ubiquitin-proteasome and lysosome-autophagy inhibitors. **(A)** Schematic of the experimental timeline showing the treatment window for bortezomib (proteasome inhibitor) or bafilomycin (autophagy inhibitor). **(B)** Immunofluorescence staining for α-actinin (red), Desmin (green), and Hoechst (blue) in CRYAB-WT-μHT and CRYAB-R120G-μHT treated with 100 nM bortezomib for 48 hours. Arrows indicate regions of sarcomeric disruption and Desmin aggregation. Zoom-in panels show that bortezomib treatment induces Desmin aggregation in both WT and CRYAB-R120G μHTs, with additional sarcomeric disorganization in CRYAB-R120G tissues. Scale bars: 50 μm (merged images), 20 μm (zoom-in). **(C)** Immunofluorescence staining for α-actinin (red), Desmin (green), and Hoechst (blue) in CRYAB-WT-μHT and CRYAB-R120G-μHT treated with 10 nM bafilomycin for 48 hours. Zoom-in panels show that bafilomycin treatment induces Desmin aggregation in CRYAB-R120G μHTs but not in CRYAB-WT μHTs, with preserved sarcomeric structure in isogenic controls. Scale bars: 50 μm (merged images), 20 μm (zoom-in). Contractility index in CRYAB-WT and CRYAB-R120G μHTs treated with **(D)** bortezomib or **(E)** bafilomycin (ratio of peak active force post-/pre-treatment). Both treatments significantly reduced contractility in CRYAB-R120G μHTs compared to isogenic controls. Data are presented as mean ± *SD*; n values represent individual μHTs pooled from ≥2 independent differentiations. *P* values by unpaired t-test, parametric; * *p* < 0.05.

To evaluate whether the CRYAB-R120G mutant μHT were similarly vulnerable to disruption of PQC through inhibition of autophagy-lysosome system or not, we treated µHTs with 10 nM Bafilomycin for 48 hours (**Fig. 4 A**). As with inhibition of the ubiquitin-proteasome system, CRYAB mutant μHT were more vulnerable than isogenic controls to inhibition of autophagy. Immunostaining confirmed this vulnerability at the structural level (**Fig. 4 C**). Desmin aggregates were present only in CRYAB-R120G µHTs following Bafilomycin treatment. Additionally, sarcomeric α-actinin organization remained relatively preserved in isogenic controls, while CRYAB-R120G µHTs showed clear disruption of sarcomeric striations. While isogenic control µHTs maintained contractile performance with minimal change (contractile index ∼1), CRYAB-R120G µHTs showed a significant drop in force production following autophagy inhibition (**Fig. 4 E**). Altogether, these observations support the hypothesis that by weakening cardiomyocyte PQC through inhibition of the heat shock protein response, mutations in CRYAB make these cells more vulnerable to inhibition of the ubiquitin-proteasome system or of the lysosome-autophagy system (53) and autophagy reactivation via TFEB (18) or ATG7 can restore proteostasis (54). Overall, cytoskeletal stress, whether mechanical, genetic, or proteotoxic, further accelerates PQC exhaustion, increasing contractile dysfunction (55, 56).

### Action potential waveform is grossly preserved in CRYAB-R120G µHTs

Given the marked deficits in contractile force and altered contractile kinetics of CRYAB-R120G mutant μHT, we sought to determine potential alternations in electrophysiology and/or calcium handling that might contribute to this pathology. We first labeled µHTs with the far-red voltage-sensitive dye BeRST-1 and recorded optical action potentials using high-speed fluorescence imaging (**Fig. S12 A**). Representative traces revealed that both CRYAB-WT and CRYAB-R120G µHTs exhibited similar action potential waveform morphology, particularly with respect to repolarization kinetics (**Fig. S12 B**). Quantitative analysis further showed no significant change in the upstroke velocity which is quantified as normalized background-corrected amplitude of the BeRST signal (ΔF/F_0_) over upstroke duration time between CRYAB-WT and CRYAB-R120G µHTs (**Fig. S12 C**). Moreover, there was no significant differences between genotypes in key repolarization metrics, including action potential durations to 30%, 50%, 80%, and 90% repolarization (APD_30_-APD_90_, respectively; **Fig. S12 D-G**). Interestingly, although total action potential duration was unchanged between genotypes (**Fig. S12 H**), the time from baseline to peak depolarization (e.g., upstroke duration), was slightly faster in CRYAB-R120G µHTs (**Fig. S12 I**), with lower background-corrected amplitude of the BeRST signal (ΔF/F_0_) (**Fig. S12 J**), leading to a similar upstroke velocity between mutant and control µHTs **(Fig. S12 C)**. Preserved action potential upstroke velocity in our study is consistent with prior electrophysiological profiling in CRYAB-R120G transgenic mice, which reported intact action potential waveforms. While our findings do not directly assess ion channel function, they align with previous evidence suggesting no alteration in Na_v_1.5 function (57).

### CRYAB-R120G µHTs exhibit impaired calcium handling

As action potential analysis suggested that impaired depolarization was unlikely to contribute to the contractility deficits of CRYAB-R120G mutant μHT, we next assessed potential dysfunction in calcium handling. To assess calcium dynamics in CRYAB-R120G µHTs, we transduced purified hiPS-CM monolayers with AAV6 vectors expressing the genetically encoded calcium indicator RGECO1.2 prior to making the μHT (**Fig. 5 A**). CRYAB-R120G tissues exhibited significantly reduced background-corrected RGECO signal intensity (ΔF/F_0_) compared to isogenic controls (**Fig. 5 B, C**). Calcium transient decay kinetics were significantly faster in CRYAB-R120G µHTs across all measured thresholds (e.g. decay from peak intensity to 75%, 50%, and 30% of peak intensity), when compared to isogenic controls (**Fig. 5 D; Fig. S13 A, B**). Furthermore, mutant μHT showed significant reduction in the upstroke velocity which is quantified as the normalized background-corrected RGECO signal intensity (ΔF/F_0_) over upstroke duration time (**Fig. 5 E)**. Moreover, similar calcium transient upstroke duration was observed in isogenic controls and mutant μHTs (**Fig. 5 F**). Importantly, the relative change in ΔF/F_0_ was significantly higher than the change in calcium transient decay, suggesting that accelerated calcium transient decay reflects overall depressed calcium intake rather than enhanced calcium handling (**Fig. 5**).

**Figure 5.**
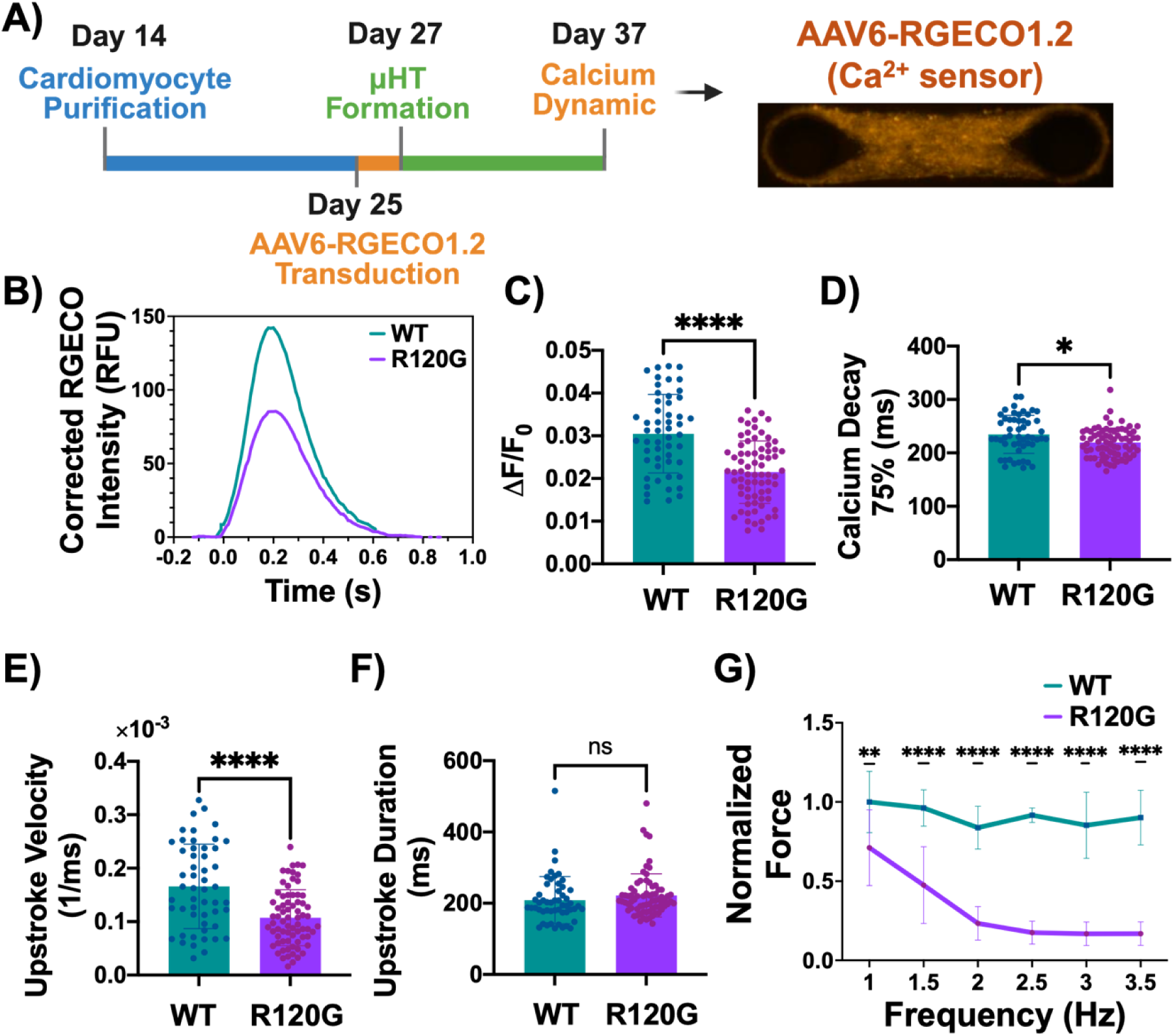
The CRYAB-R120G mutation impairs calcium handling in 3D μHTs. (A-F) Calcium measurements using AAV6-RGECO1.2. **(A)** Experimental timeline and representative image of μHT expressing RGECO1.2 (genetically encoded Ca^2+^ sensor). **(B)** Representative corrected RGECO1.2 intensity traces from CRYAB-WT and CRYAB-R120G μHTs. Quantification of **(C)** background-corrected RGECO signal intensity (ΔF/F_0_) and **(D)** Ca^2+^ decay time to 75% of peak. CRYAB-R120G μHTs show significantly reduced Ca^2+^ transient amplitude and faster decay kinetics. Quantification of **(E)** Ca^2+^ upstroke velocity measured by normalized background-corrected RGECO signal intensity (ΔF/F_0_) over upstroke duration time and **(F)** Ca^2+^ upstroke duration time (UPD). Data are presented as mean ± *SD*; n values represent individual μHT pooled from ≥3 independent differentiations. *P* values by unpaired parametric Student’s t-test for ΔF/F_0_ and Mann–Whitney test for non-parametric data (upstroke velocity, upstroke duration time, and Ca^2+^ decay 75%); **** *p* < 0.0001, * *p* < 0.05, ns = not significant. **(G)** Normalized force–frequency relationship in CRYAB-WT and CRYAB-R120G μHTs paced from 1 to 3.5 Hz, showing a pronounced negative force–frequency response in CRYAB-R120G μHTs. Values are normalized to the mean WT peak active force at 1 Hz pacing. Data are presented as mean ± *SD*; n values represent individual μHT pooled from one differentiation. Data were analyzed by two-way ANOVA, followed by Holm–Šidák’s multiple comparison test.

Furthermore, reduced Ca^2+^ intake, quantified via lower values for background-corrected RGECO signal (ΔF/F_0_) **(Fig. 5 C),** is consistent with the hypothesis that the CRYAB-R120G mutation diminishes sarcoplasmic reticulum function, leading to impaired calcium-induced calcium release (58). Depressed sarcoplasmic reticulum function would be expected to manifest in exaggerated contractile deficits at higher pacing frequencies because of inhibited calcium cycling (58). We thus assessed the force-frequency response of the μHT. Isogenic control tissues exhibited a largely flat force-frequency response up to a pacing frequency of 3.5 Hz (**Fig. 5 G**). This is consistent with prior reports of a moderate degree of physiological maturation of hiPS-CM in engineered heart tissues (59, 60). However, CRYAB-R120G mutant μHT exhibited a markedly negative force-frequency response (**Fig. 5 G)**, consistent with the hypothesis that depressed calcium transient amplitude is linked to a dysfunctional sarcoplasmic reticulum in these cells. Altogether, these findings suggest that calcium handling dysfunction plays a key role in the hypocontratile phenotype of the CRYAB-R120G mutant μHT.

### Itacitinib treatment reduces STAT1 activation and partially rescues contractile deficits in CRYAB-R120G µHTs

In prior work, an unbiased screen performed in neonatal rat ventricular cardiomyocytes identified JAK/STAT signaling in the generation of proteotoxic aggregates in the setting of CRYAB-R120G overexpression (38). Subsequent investigation suggested that JAK/STAT signaling via JAK1 inhibitors, including Itacitinib, could effectively remove such aggregates (61, 62). We thus tested the potential of JAK1 inhibition via Itacitinib as a candidate treatment for DRM. μHTs were exposed continuously to Itacitinib for one week (**Fig. 6 A**). Following treatment, we fractionated the lysates of μHT to separately assess the soluble (cytoplasmic) and insoluble (aggregate and cytoskeletal) fractions to assess STAT1 signaling (**Fig. 6 B)**, along with CRYAB expression and portioning into detergent-insoluble aggregates, by Western blot.

**Figure 6.**
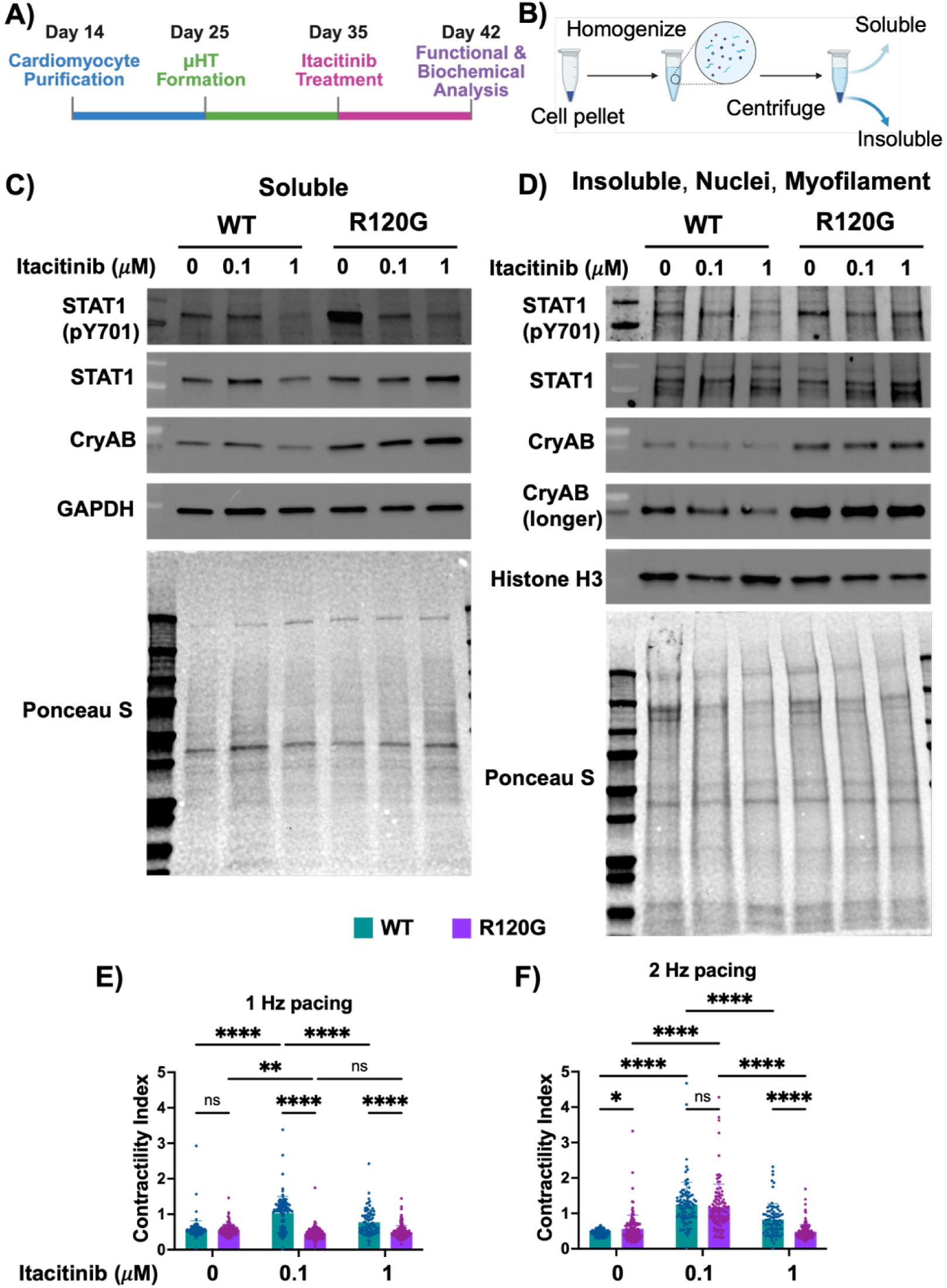
Itacitinib reduces STAT1 activation but does not fully rescue contractility deficit in CRYAB-R120G μHTs. **(A)** Schematic of the experimental timeline for treating μHTs with Itacitinib. hiPS-CMs were metabolically purified (D14–D25) after differentiation, seeded into 3D μHT devices (D25–D35), and treated with 0.1 and 1 μM Itacitinib and vehicle control from D35–D42 before analysis. **(B)** μHTs were lysed using RIPA buffer with 1% NP-40 into soluble and insoluble fractions. **(C)** Representative immunoblots of soluble fractions showing dose-dependent reduction in phosphorylated STAT1 (pY701) in both CRYAB-WT and CRYAB-R120G μHTs following treatment with 0.1 or 1 μM Itacitinib, with total STAT1 unchanged. Soluble CRYAB levels were higher in CRYAB-R120G μHTs compared to CRYAB-WT across all conditions. GAPDH was used as a loading control; Ponceau S staining shown as reference. **(D)** Representative immunoblots of insoluble fraction containing aggregates/nuclear/myofilament fractions showing dose-dependent reduction of phosphorylated STAT1 with Itacitinib treatment, while total STAT1 levels remained unchanged. CRYAB accumulated at higher levels in insoluble fractions of CRYAB-R120G μHTs, regardless of treatment. Histone H3 was used as a loading control; Ponceau S staining shown as reference. Blots are representative of 2 individual differentiation batches. Contractility index (ratio of peak active force post-/pre-treatment at the same μHT) treated with vehicle control, 0.1 μM, and 1 μM Itacitinib paced at **(E)** 1 Hz and **(F)** 2 Hz frequency. 0.1 μM Itacitinib partially rescued contractility at 2 Hz; however, 1 μM Itacitinib worsened contractility. Data are presented as mean ± *SD*; n values represent individual μHTs pooled from ≥2 independent differentiations. Data were analyzed by two-way ANOVA, followed by Holm–Šidák’s multiple comparison test. **** *p* < 0.0001, *** *p* < 0.001, ** *p* < 0.01, * *p* < 0.05, ns = not significant.

In the soluble fraction (**Fig. 6 C)**, treatment with Itacitinib led to a dose-dependent reduction in phosphorylated STAT1 (pY701) in both isogenic controls and CRYAB-R120G tissues, confirming effective inhibition of JAK/STAT signaling. Total STAT1 levels were higher in mutant µHTs, however, overall STAT1 levels remained unchanged with drug treatment. Soluble CRYAB levels also appeared higher in mutant µHTs than isogenic controls, consistent with our immunofluorescence data (**Fig. 2 E, F**).

In the insoluble lysate fraction, which includes aggregated, nuclear, and myofilament proteins (**Fig. 6 D**), phosphorylated STAT1 was also decreased in a dose-dependent manner for both isogenic controls and mutant µHTs. Total STAT1 levels also remained unchanged across different treatment conditions. Moreover, CRYAB level was higher in the insoluble fraction of CRYAB-R120G tissues regardless of treatment compared to isogenic controls (**Fig. 6 D)**.

To assess the functional consequences of JAK1 inhibition, contractile performance was measured before and after treatment. Based on our finding that the force-frequency response is especially impaired in DRM-μHT (**Fig. 5 G**), we assessed contractile performance at 1 Hz and 2 Hz. Strikingly, in isogenic control (CRYAB-WT) μHT, Itacitinib treatment significantly improved peak active force (**Fig. S14 A**) at both 0.1 µM and 1 µM doses under both pacing conditions; with improved contractility index (ratio of peak active force before and after drug exposure) at both frequencies (**Fig. 6 E, F**). In contrast, CRYAB-R120G μHTs demonstrated recovery of contractility at 2 Hz following treatment with 0.1 µM Itacitinib without significant improvement at 1 Hz (**Fig. S14 A)**. While absolute contractility at this pacing frequency did not match isogenic control μHT treated with the same drug dose (0.1 μM), it was not significantly different from isogenic control tissues treated with vehicle (**Fig. S14 A**). 0.1 µM Itacitinib also improved the relaxation velocity in isogenic controls at both 1 and 2 Hz pacing frequency (**Fig. S15 A, B**). However, drug-treated CRYAB mutant μHT (0.1 µM) failed to match the relaxation velocity of vehicle-treated isogenic controls (**Fig. S14 B**). These results suggest that while JAK/STAT inhibition enhances contractile function in μHT, regardless of genotype, it partially rescues contractility in the context of CRYAB-R120G induced proteotoxicity. Interestingly, higher doses of Itacitinib did not further enhance contractility and instead appeared to impair function, suggesting that JAK/STAT signaling may have dual role. Prior work on engineered heart tissues has shown that sustained IFN-γ exposure drives STAT1 activation, leading to loss of contractile force and sarcomere disarray; and these detrimental effects are prevented by JAK inhibition (63). However, *in vivo* studies of pressure overload induced hypertrophy have revealed a protective role of STAT1 when stabilized by the deubiquitinase USP13. Specifically, USP13 enhances STAT1 deubiquitination and stability, promoting its transcriptional activity, improving mitochondrial function, and attenuating TAC- and AngII-induced hypertrophy (64). Thus, excessive STAT1 activation under inflammatory stress (e.g., IFN-γ) is maladaptive, while basal STAT1 stabilization under biomechanical stress can be cardioprotective. This dual role likely explains why higher-dose JAK/STAT inhibition in our model compromises function, highlighting the delicate balance between pathological and adaptive STAT1 signaling.

## Discussion

PQC systems are essential for maintaining cardiac function by ensuring the integrity of sarcomeric and cytoskeletal proteins (18, 65, 66). In this study, we developed, to our knowledge, the first *in vitro* human model of DRM using hiPS-CMs harboring a homozygous CRYAB-R120G knock-in mutation expressed at endogenous levels, which are subjected to physiologically relevant mechanical loading. Unlike previous DRM models that relied on supraphysiological overexpression of mutant CRYAB (27), our genome-edited platform preserves physiological CRYAB-client protein expression balance, which enable a more accurate assessment of disease mechanisms and potential underlying protein-protein interactions. This model recapitulates hallmark DRM phenotypes, including Desmin aggregation, cellular hypertrophy, contractile dysfunction, altered calcium handling, and heightened vulnerability to proteostasis disruption within human cardiomyocytes. We demonstrate that JAK/STAT pathway inhibition, previously proposed as a candidate for DRM therapy (61, 62), led to modest improvements in contractile function.

Our findings further support the role of CRYAB in modulating Desmin filament structure; prior studies indicate this occurs through direct, phosphorylation-dependent CRYAB-Desmin binding (67–69). In CRYAB-R120G µHTs, increased Desmin aggregation and sarcomeric disorganization were observed, consistent with electron-dense aggregates in previous reports (6, 28, 70). In those prior studies, these aggregates, with CRYAB at the core and Desmin at the periphery, displaced Desmin from Z-discs and intercalated discs into cytosolic inclusions, which also contained α-actinin and ubiquitin (18, 21, 71), indicative of collapsed sarcomere structure. Functionally, CRYAB-R120G µHTs showed reduced peak active force and slower contraction-relaxation kinetics, paralleling prior evidence that proteotoxic sarcomeric protein aggregates disrupts Z-disc alignment and force transmission, therefore driving contractile deficits in DRM (28, 71, 72) . In addition, these prior studies report mitochondrial abnormalities, such as reduced Complex I and IV activity, swelling, cristae loss, and impaired ATP production, which could also inhibit contractility (71, 73, 74).

While prior studies have emphasized roles for direct sarcomeric and mitochondrial damage in CRYAB-DRM associated contractile dysfunction (28, 71–74), our findings suggest a significant defect in calcium handling in CRYAB-R120G mutant µHTs (**Fig. 5**). This was supported not only by direct analysis of Ca^2+^ handling, but also by a markedly negative force-frequency response in the mutant tissues. Together, these findings implicate insufficient sarcoplasmic reticulum calcium loading, reflecting impaired refilling due to RyR2, SERCA2, or PLN dysfunction (75). Prior reports indicate that loss of chaperone function in CRYAB-R120G mutants can lead to destabilization of SERCA2 (76), reducing the sarcoplasmic reserve of calcium (58). This effect likely reflects chronic stress on the sarcoplasmic and endoplasmic reticulum, as aggregation of misfolded CRYAB-R120G proteins exacerbates sarcoplasmic and/or endoplasmic reticulum stress and impairs calcium-handling capacity (76, 77).

Therapeutically, JAK1 inhibition with Itacitinib partially restored contractile performance, consistent with expectations based on prior work showing that CRYAB-R120G aggregation is largely rescued by JAK1 inhibition (61, 62). Indeed, clinically approved JAK1 inhibitors enhance ubiquitin-proteasome aggregate clearance via muscle-specific E3 ligases such as Asb2 (38, 78), highlighting JAK1 as a promising target for DRM therapy. However, our finding indicates that JAK1 inhibition only partially rescued contractility of CRYAB-R120G µHTs, with the effect more pronounced at higher pacing frequencies, while evidence also suggests that doses higher than 0.1 µM have diminished physiological benefit and do not rescue CRYAB-R120G partitioning into aggregates. This suggests the potential for a more multifaceted role for JAK/STAT signaling in DRM suggesting that a complete shutdown of JAK/STAT signaling may be deleterious, whereas finely tuned modulation is protective (79, 80). Prior studies highlight the dual nature of JAK/STAT signaling. In ischemia–reperfusion models, excessive activation of JAK/STAT exacerbates injury, whereas its inhibition reduces infarct size and apoptosis (81). Similarly, Pan *et al.* showed that mechanical stretch rapidly activates JAK/STAT in cardiomyocytes via gp130 cytokines, linking physiological stress to hypertrophic adaptation (82). Moreover, chronic or excessive pathway activation has been consistently associated with maladaptive remodeling, hypertrophy, and fibrosis (80). In contrast, consistent with the potential for JAK/STAT signaling to play a beneficial role in cardiomyocytes, especially in the setting of mechanical stress, recent work suggests that loss of STAT1 is deleterious in cardiac hypertrophy provoked by transaortic constriction *in vivo* (64), highlighting the context-dependent nature of this pathway.

Our findings on the importance of JAK/STAT signaling in μHT, and a potential need for basal, but not excessive, levels of these signals, also align with work in engineered human myocardium. Zhan *et al.* reported that IFN-γ exposure, which strongly upregulates JAK1/STAT1, caused sarcomeric protein loss and contractile decline, effects that were fully rescued by JAK inhibitors (63). In contrast, Ho *et al.* reported that IFN-γ–induced STAT1 activation promoted maturation of hESC-derived cardiomyocytes, enhancing calcium handling, sarcomere alignment, and metabolic switching to fatty acid oxidation which were reversed by ruxolitinib (83, 84). Taken together, these studies and our results suggest that modest JAK/STAT activity, is essential for homeostasis, maturation, and stress adaptation. However, chronic overactivation or complete pharmacological suppression can both impair cardiac function. The diminished rescue we observed at higher Itacitinib doses may therefore reflect the loss of necessary adaptive pSTAT1.

Our finding that Desmin expression was preferentially induced in 3D µHT is consistent with reports that 3D cardiac aggregates show higher Desmin expression than 2D cultures (85). Prior work from our team (46, 50) and others (86–88) highlight that hiPS-CM change their biology in physiologically relevant culture systems, which can facilitate disease modeling. Desmin is the major muscle-specific intermediate filament protein in cardiac, skeletal, and smooth muscle. It forms a 3D scaffold around Z-disks, linking myofibrils to the sarcolemma, mitochondria, nuclei, and cell junctions. Although not essential for early development (as reflected in our finding that Desmin is absent in immature cardiomyocytes) it becomes indispensable postnatally for maintaining structural integrity and function (66).

The mechanical function of Desmin, coupled with the mechanical stresses that hiPS-CM experience within μHT, suggest that mechanical stress is key to Desmin expression and/or stability. Desmosomes are likely to be more prevalent in 3D, and Desmin intermediate filaments directly interface with desmosomes to bear mechanical stress during coordinated contraction. Recent work shows that desmosome–intermediate filament systems actively facilitate mechanotransduction at adherens junctions by amplifying tension-sensitive RhoA signaling (89). Moreover, Desmin has been shown to integrate 3D mechanical properties of muscle by coupling transverse and longitudinal stress and contributing to load-dependent stiffness and viscoelasticity, suggesting that its expression is sensitive to mechanical load (90). Thus, the induction of Desmin expression in 3D μHT may represent an adaptive response to the stress-bearing and mechanosignaling role of desmosomes, reinforced by ECM-mediated signaling, whereas in 2D cultures, components of cell-cell junctions, including desmosomes, would be expected to experience less mechanical loading, thereby requiring less reinforcement from Desmin intermediate filaments for structural integrity (85, 91, 92).

In summary, our CRYAB-R120G µHT model recapitulates the complex interplay between Desmin aggregation, PQC exhaustion, altered calcium handling, and inflammation. This positions DRM as a prototypical model for studying PQC failure and sarcomere degeneration, with our system offering a versatile platform for dissecting mechanisms and screening therapies in both inherited and acquired cardiomyopathies.

Compared to prior experimental systems for modeling PQC-associated DRM, our model has several advantages. First, previous DRM models using transgenic overexpression, which often required excessive CRYAB levels (28), may have induced artificial protein-protein interactions (93), whereas our model maintains endogenous gene expression dosage. Second, 3D µHT culture promotes cardiomyocyte structural maturation, which includes Desmin expression, similar to findings in previously described, more complex, macroscale engineered heart tissue models (35), albeit with a requirement for ∼20-fold fewer cells per construct, which offers advantages for screening applications. Third, our data align with prior studies showing that Desmin functions as a cytoskeletal scaffold critical for cardiomyocyte structural integrity (65, 66), and that its organization is reinforced under mechanical loading in 3D systems similar to dyn-EHT (35). Considering that chemotherapy agents such as doxorubicin and bortezomib disrupt PQC by impairing proteasomal degradation and autophagic flux (94), our model of familial DRM may also be useful in future studies to identify potential treatments for both genetic and drug-induced cardiomyopathies involving proteotoxic stress.

Limitations of our μHT model include the short culture duration, and use of a homozygous genotype rather than the more common heterozygous patient genotype (6). Future studies incorporating single-cell omics, mitochondrial stress assessment, and heterozygous patient-derived lines will likely enhance translational relevance, while live imaging of aggregation could reveal dynamic mechanisms of disease progression.

## Methods

### Human iPSC Culture and Differentiation

Human induced pluripotent stem cells with mKate-tagged α-actinin were previously described (95). These cells were targeted with CRISPR-Cas9 at the Genome Editing and Stem Cell Center at Washington University School of Medicine to generate homozygous knock in lines to change codon for expression of arginine to glycine at position 120. We thank the Genome Engineering & Stem Cell Center (GESC@MGI) at Washington University in St. Louis for reagent validation services. The mutation was validated by next generation sequencing and resulting R120G mutation in hiPSC. Isogenic controls (CRYAB-WT-hiPSC) and mutant (CRYAB-R120G-hiPSC) were cultured in MTesR1 media (Fisher Scientific; NC1274780) and gradually adapted to StemFlex media (Life Technologies; A3349401). During passaging and differentiation, hiPSCs were passaged as single cells using Gentle Dissociation Reagent (PBS with 1.8g/L NaCl and 0.5mM EDTA; (96)), and replated into media supplemented with 10 μM Y27632 (Biogem; 1293823) onto plates coated with Geltrex (Life Technology; A1413302) at a concentration of 18.75 μg/cm^2^. For differentiation, hiPSCs were subjected to timed control over Wnt signaling using small molecules (34, 50). Briefly, hiPSCs were seeded at a density of 40,000 cells/cm^2^ onto Geltrex (37.5 μg/cm^2^) and expanded for 3 days until confluent in MTeSR or StemFlex media. Next, media was changed to RPMI1640 (Life Technologies; 11875093) supplemented with 150 μg/ml ascorbic acid (Fisher Scientific; NC0602549) and 2% B27 without insulin (RPMI-I) (Life Technologies; A1895601) containing 6 μM CHIR99021 (Biogem; 2520691) (differentiation day 0). After 2 days, media was changed to RPMI-I containing 5 μM IWP2 (Biogem; 6866167). After 2 additional days, media was changed to RPMI-I. On day 6, media was changed to RPMI1640 supplemented with 2% B27 (Life Technologies; 17504044) (RPMI-C). Thereafter, culture was fed every 2 days with RPMI-C. Beating cardiomyocytes were typically observed by differentiation day 8. On differentiation day 14, beating cardiomyocyte monolayer were gently dissociated using 10X TrypLE Select Enzyme (Gibco; A1217701). The single cells were then replated onto Geltrex-coated tissue culture plastic (37.5 μg/cm^2^) at a density of 300,000 cells/cm^2^ and cultured in RPMI-C media supplemented with 20% fetal bovine serum (FBS; Fisher Scientific; FB12999102) and 10 μM Y27632 for 24 hours. At day 15, the media was changed to a lactate-based metabolic selection media (RPMI1640 without glucose (Millipore Sigma; R1383) supplemented with 4mM lactate (Fisher Scientific; AAL1450014), 1% Non-Essential Amino Acids (Fisher Scientific; 11140050), and 1% Glutamax (Fisher Scientific; 35050061). On day 18, the media was replaced with fresh metabolic selection media. After 6 days culture in metabolic selection media, from day 21 to 24, the media was gradually changed back to RPMI-C for post-selection recovery. At differentiation days 24-26, purified hiPS-cardiomyocytes (**Fig. S11 A**) were used for making μHT (**Fig. 1 A**).

### Monolayer Study

Purified cells were dissociated and seeded onto Geltrex-coated Permanox chamber slides (Fisher Scientific; 1256522) at density of 50,000 cells/cm^2^. The replating media is RPMI-C with 10 μM Y27632 and 10% FBS. For maturation media experiment, the media was changed to maturation media (MM) consist of 125 μM palmitic acid (Fisher Scientific; NC1980206), 125 μM Linoleic acid, 125 μM oleic acid (Millipore Sigma; L9655), and 5 μM GW0742 (MedChemExpress; HY-13928) as PPAR-δ agonist in RPMI-without glucose supplemented with 2.8 mM Glucose, and 10 mM Galactose (41, 42). The media was changed to either fresh MM or fresh standard media (SM) (RPMI-C) every 2 days for 2 weeks.

### Device Design

The mold for the platform was designed in Autodesk Fusion 360 to fabricate 48 microwells, each 1.25 mm deep, containing two tapered posts 1 mm in height, with a 270 µm base diameter and a – 2° taper. The posts are capped to retain the tissues on their top, with the caps chamfered at 1° on both the top and bottom edges while the central portion remains straight (total cap height: 250 µm). The distance between the two posts is kept as 1850 µm. The finalized designs were exported as .stl files and uploaded to PreForm software for slicing, then printed on a Form 3 printer with a 50 µm layer thickness (**Fig. S2 A, B**).

### Device Fabrication

Device fabrication was explained previously (44, 97). Briefly, PDMS posts were fabricated using Hydrogel Assisted Stereolithographic Elastomer prototyping. After getting the master print from Form 3 printer, devices were washed in dPBS and dried. To create the inverse-shape of the posts, agar (Fisher Scientific; 1.5%wt/v in tap water) was boiled. Triton-X-100 was mixed to a final concentration of 0.2% during the time it took for the agar to cool below the glass transition temperature of the 3D prints. After reaching 60°C, the agar/Triton-X-100 mixture was poured onto the 3D printed part, cooled down, and cross linked for 30 minutes at 4°C. Next, the 3D print was gently removed from the agar, leaving the agar mold as an inverse shape. Sylgard 184 mixture was prepared by mixing the crosslinker and base with 1:10 ratio and degassed under vacuum chamber. In parallel, Sylgard 527 mixture was prepared by mixing part A and part B with 1:1 ratio, then degassed in the same way. Afterward, to make the stiff posts with elastic modulus of 3000 kPa and theoretical bending stiffness of 1.96 N/m, pure Sylgard 184 was poured into the agar mold. In parallel, to make soft posts, Sylgard 184 mixture was mixed with Sylgard 527 mixture with 1:4 ratio, to yield PDMS with elastic modulus of 640 kPa and theoretical bending stiffness of 0.42 N/m. After pouring the PDMS mixture into the agar mold, it was degassed and kept at 37°C oven overnight. The next day, PDMS devices were removed from agar and underwent final crosslinking at 60°C overnight.

In the meantime, we fabricated polycarbonate inserts to contain the PDMS devices. Polycarbonate inserts were created by thermoforming against a tooling mold 3D-printed in high-temperature resin on a Phrozen Digital Light Printing (DLP) 3D printer as described previously (98). After forming the polycarbonate inserts, PDMS post devices were attached to the inserts using Sylgard 184 as a glue, and the devices were then autoclaved and stored under aseptic conditions until ready for cell seeding to form μHT.

### Surface Modification

To reduce the cell attachment to the sterilized PDMS devices, we submerged the devices in 1% wt/v Pluronic F127 for at least 2 hours at room temperature, followed by 3 washes (five minutes each) in sterile dPBS.

### iPSC-μHT Formation and Optimization

Purified cardiomyocytes were gently dissociated using 10X TrypLE Select Enzyme (for up to 15 minutes). In parallel, human primary ventricular cardiac fibroblast (Lonza; CC-2904) were dissociated using 0.25% trypsin EDTA (Fisher Scientific; 25200072) for 5 minutes. A mixture of 5% primary cardiac fibroblast and 95% of hiPS-CMs, at a total cell density of 2×10^7^ cells/ml were encapsulated into 1.5 mg/ml collagen I rat tail (Fisher Scientific; A1048301) and 10% v/v Geltrex. 4 μL of this mixture (80,000 cells) was seeded into each 2-post μHT-forming device (**Fig. S2 C**). The seeded devices were next incubated at 37°C with 5% CO2 for 20 minutes to crosslink collagen, before the entire culture well was filled with μHT culture media (KnockOut DMEM (Life Technologies; 10829018) supplemented with 20% FBS, 1% Glutamax, 1% Non-essential Amino Acids, 1% Pen Strep (Fisher Scientific; 15-140-122), 150 μg/ml ascorbic acid and 10 μM Y27632). Tissue compaction was typically observed within 48 hours (**Fig. S2 C, D**). After 2 days, media was changed to KnockOut DMEM supplemented with 1% KnockOut Serum Replacement (Fisher Scientific; 10828010), 1% Glutamax, 1% Non-essential Amino Acids, 1% Pen Strep, and 150 μg/ml ascorbic acid. The media was refreshed every 2 days until day 10 in tissues.

### Immunostaining

μHTs and monolayers were incubated in 100 mM KCl until cessation of spontaneous beating, and then step fixed with 1%, 2%, 3% and 4% paraformaldehyde at 4°C, with each fixation step at least 1 hour, and washed in dPBS. These tissues were subsequently used for immunostaining for sarcomeric α-actinin (Millipore Sigma; A7811), Desmin (Cell Signaling; 5332), and CRYAB (Enzo Life Science; ADI-SPA-223). Wheat germ agglutinin (WGA) (Fisher; W11261) was used for staining cell membranes, and cell nuclei were stained with Hoechst 33342. Whole mount tissue or monolayer stained were mounted in ProLong Gold (99) (Fisher Scientific; P36930) and scanned using a Fluoview FB1200 confocal microscope (Olympus, Tokyo, Japan). In-house structural analysis codes are available here: (https://github.com/huebschlab/CardioSpatialAnalysis).

### AAV6-driven-ChR2(H134R) and/or RGECO1.2 Constructs and Transduction

pAAV-CMV-CHR2-eYFP was from Vectorbuilder (VB240118-1580mnz). pAAV-CMV backbone was generously provided by Dr. Brent French at University of Virginia, Charlottesville, VA. CMV-RGECO1.2 was a gift from Robert Campbell (Addgene plasmid #45494; http://n2t.net/addgene:45494; RRID:Addgene_45494; (100)).

To generate pAAV-CMV-ChR2(H134R)-eYFP-P2A-RGECO1.2 construct, fragments encoding ChR2(H134R)-eYFP and p2a-RGECO1.2 were PCR-amplified, restriction-digested, and ligated into the pAAV-CMV backbone using T4 DNA ligase. Adeno-associated virus serotype 6 (AAV6) particles coding for ChR2(H134R)-eYFP or ChR2(H134R)-eYFP-p2a-RGECO1.2 driven by CMV promoter were generated by the Hope Center viral vectors core.

For AAV transduction, hiPS-CM monolayers were transduced following metabolic purification at day 25 (MOI: 5 × 10^3^, yielding ∼100% transduction efficiency). Cells were exposed for 6 hours in infection medium (RPMI-C, 2% FBS, 1% Glutamax, 0.1% penicillin-streptomycin, 1% HEPES), then recovered in culture medium (RPMI-C, 10% FBS, 1% Glutamax, 0.1% penicillin-streptomycin, 1% HEPES) prior to μHT formation. Resulting μHT robustly expressed the vector, exhibited RGECO1.2 based Ca^2+^ transient, and could be optogenetically paced by blue light for > 30 days.

### High-speed Epifluorescence Imaging

Physiology of hiPSC-μHT at tissue day 10 (corresponding to day 35 of hiPS-cardiomyocytes differentiation) was analyzed using a high-speed imaging system (Eclipse Ts2R, Nikon; Tokyo, Japan) equipped with a digital CMOS camera (Hamamatsu ORCA-Flash4.0 V2; Japan). The imaging stage was set to a constant 37°C temperature using a thermal plate (Tokai Hit; Shizuoka, Japan). Field pacing of tissue (1 Hz, 20 msec bipolar pulses, <20V) during imaging was achieved using graphite electrodes (MyoPacer, Ion Optix, USA). 1 Hz field pacing was chosen because it is slightly above the average spontaneous beating rate of the cardiac tissues. In parallel, instead of field pacing with graphite electrodes, we used AAV6-ChR2 at MOI of 5,000 to pace the tissues using GFP blue light. We also used AAV6-ChR2-RGECO1.2 to measure the calcium dynamic with Texas red light.

### Contractility Characterization

Open source Beat Profiler software (101) was used to quantify tissue contractility. As an input to this software, the theoretical bending stiffness of our μHT posts were determined by finite element model using Ansys software. For soft posts (elastic modulus = 640 kPa), the bending stiffness was 0.42 N/m, corresponding to a 98 μN force required for 234 μm deflection. For stiff posts (elastic modulus = 3000 kPa), the bending stiffness was 1.96 N/m, corresponding to a 98 μN force for 50 μm deflection.

### Action Potential and Calcium Dynamic Measurements

Membrane potential was measured by labeling μHTs with 1 μM BeRST-1 voltage sensitive dye for 5 hours in phenol red free RPMI-C. Action potential waveforms were captured by imaging in the Cy5 channel. Calcium transients were captured by imaging RGECO1.2 in the Texas red channel. BeRST-1 and RGECO1.2 videos were analyzed via custom, open-source MATLAB code (available at: https://huebschlab.wustl.edu/resources-2/; (102)); intensity profile and kinetics parameters were automatically extracted. For calcium transient dynamics, initial baseline to peak of the intensity was measured as UPD, times from the peak to 30, 50 and 75% decay of intensity was characterized as decay30, decay50, and decay70. For action potential waveforms, initial baseline to peak intensity was measured as UPD, and the time from action potential initiation to 30% decay from peak intensity (APD_30_), along with the time from action potential initiation to 80% or 90% decay from peak intensity (APD_80_ and APD_90_, respectively), were measured.

### Pharmacology Studies

The effect of sarcomeric PQC pathways for pathologic contractility in CRYAB-R120G induced cardiomyopathy was investigated using pharmacological probes. Bortezomib was chosen as ubiquitin-proteasome inhibitor (100 nM), and Bafilomycin (10 nM) as lysosome-autophagy inhibitor. Drugs were first diluted in dimethyl sulfoxide (DMSO), then serial diluted in tissue culture media with supplements. Before exposure to drug the tissues were imaged for contractility measurement, then these drugs were used for 48 hours at 37°C with 5% CO_2_ then imaged for contractility again to evaluate the chronic exposure for protein turnover. Finally, we calculated the contractility index as the ratio of the peak active force after exposing to drugs over before exposing to drug followed by fixation and immunostaining.

To study the potential impact of JAK/STAT inhibition as a therapy for DRM, we treated µHTs with 0.1 µM and 1 µM Itacitinib, a JAK1 inhibitor, for 1 week to assess the role of inflammatory signaling pathways, specifically examining pSTAT1 (Tyr701) and STAT1 levels. Fresh drug was added every day.

### Immunoblotting

Soluble and insoluble protein fractions were isolated from µHTs by sequential extraction. Tissues were first lysed in RIPA buffer (Cell Signaling; 9806) supplemented with 1% NP-40 to extract the soluble protein fraction, collected as the supernatant following centrifugation. The remaining pellet was then solubilized in 9 M urea to obtain the insoluble protein fraction. Immunoblotting was performed as previously described (18). Specific antibodies are as follows: CRYAB (Enzo Life Science; ADI-SPA-223), pSTAT1(Tyr701; Cell Signaling; 58D6), STAT1 (Thermo Fisher; AHO0832), GAPDH (Abcam; ab22555), and Histone 3 (Cell Signaling; 9715).

### Quantification and Statistical Analysis

GraphPad 9.3.1 was used for statistics and data visualization. Data are presented with mean ± standard deviation (*SD*) and are pooled from μHT (or 2D monolayers) from at least 3 independent differentiation batches. For quantitative analysis, data were first assessed for normality using Normality and Lognormality tests. If datasets were normally distributed, comparisons between two groups were performed using a parametric Student’s t-test, and comparisons among multiple groups were analyzed using parametric one-way ANOVA followed by Holm–Šidák post hoc test for multiple comparisons. If datasets were not normally distributed, a non-parametric Mann–Whitney U test was used for two-group comparisons, and one-way ANOVA with Dunn’s correction was applied for multiple group comparisons. When two independent variables were compared, data were analyzed using two-way ANOVA followed by Holm–Šidák post hoc test. All statistical analyses were performed as two-tailed tests unless otherwise specified.

### Sources of Funding

A.D. was supported by grants from the National Institutes of Health (HL107594) and the Department of Veterans Affairs (I01BX005065, I01BX005981). D.R.R. is supported from grants from the National Institutes of Health (T32 HL007081 and 1K08HL163469). NH was supported by grants from National Institutes of Health (HL159094) and National Science Foundation CAREER (2338931). G.R. and H.Y.C were supported by McDonnell Academy Fellowship. J.S.P. was supported by T32GM007200

## Disclosure

N.H. reports consulting for Organos, Inc. (Moraga, CA), which did not affect the current study. A.D. reports consulting for clinical trials with Clario (previously ERT/Biomedical systems); and serving on the scientific advisory board for Dewpoint Therapeutics, which did not affect the current study.

