## Supporting Information for "A Micro-Engineered Heart Tissue Model of Desmin-related Cardiomyopathy Caused by Mutant αB Crystallin"

**Supplementary Method:**

### 1.1 Cell Segmentation with CellPose (Cyto2) model:

Nuclei and α-actinin stained images were paired and organized prior to segmentation, ensuring that only correctly matched image sets were processed. Images were converted to float32, normalized to a [0, 1] range, and stacked into two-channel arrays with α-actinin assigned as the first channel to ensure accurate interpretation by CellPose. Segmentation was performed using the cyto2 model in 2D mode with an initial diameter of 100 pixels. Resulting masks were used to extract cellular morphology features including area, elongation, eccentricity, circularity. Measurements were exported to a single CSV file, and false-color overlays were generated with region IDs to enable visual quality control. Two strategies were applied to optimize segmentation. First, direct application of cyto2 with diameter adjustments was used to improve agreement between predicted and observed α-actinin cell sizes. Second, the model was fine-tuned using wheat germ agglutinin (WGA) images, which provide well-defined membrane boundaries, thereby improving segmentation accuracy on α-actinin datasets.

### 1.2 Tissue segmentation with nuclei:

Nuclei were segmented from Hoechst stains using Gaussian filtering, Otsu thresholding, and watershed separation. Small debris and border-touching nuclei were excluded, yielding clean nuclear seeds. Approximate whole-cell regions were then generated by expanding each nucleus outward until neighboring boundaries met, producing a Voronoi-like tessellation of the tissue.

From these labeled boundaries, cellular morphology features (area, elongation, eccentricity, and circularity) were extracted. Desmin intensity was quantified on a per-cell basis, both as mean and total intensity. To assess Desmin positivity, an Otsu threshold was applied to the

Desmin channel, and the marker-positive fraction was calculated for each cell. Outputs included scatter and heatmaps of Desmin-positive area as well as false-color overlays of cell boundaries.

**1.3 Identification of Desmin aggregation:**

For Desmin aggregation analysis, nuclei were first segmented from Hoechst staining to define cell boundaries (**Fig. 1E; Supplemental Video 2)**, and Desmin images were processed with SarcAsM software (1) to identify Z-band-like structures, corresponding to Z-disc localized Desmin. These "Z-band-like structures" were used to create a binary mask of "organized Desmin". Next, to avoid falsely counting pixels with weak Desmin signal as having disorganized Desmin, *intensity* values were used directly to generate a second binary mask, corresponding to pixels occupying the $75^{th}$ percentile (or higher) of Desmin signal intensity. Cells with fewer than 100 Desmin-positive pixels were excluded. Within each cell, we then counted 1) the total number of pixels having "Z-band-like" Desmin structure, and 2) the total number of pixels having appreciable Desmin signal. Pixels that had appreciable Desmin signal but no Z-band-like structure were considered aggregated. For quantification, we summed data from all cells within each ROI and selected two representative ROIs per μHT, from a total of 20 μHTs pooled from three independent hiPS-CM differentiation batches. This formal, automated analysis produced results similar to what we observed when a trained user manually selected "aggregated" vs. "organized" areas of Desmin staining.

**1.4 Analysis of spatial distribution of Desmin-positive cardiomyocytes in μHT:**

To assess whether Desmin expression in μHTs occurred in a clustered or random spatial pattern, nuclei were first segmented from Hoechst staining to define cell boundaries (**Fig. 1E; Supplemental Video 2)**, and Desmin intensity was measured per cell. Cells above a set intensity threshold were classified as Desmin-positive. An adjacency graph was then constructed to connect

Desmin-positive cells that shared a boundary with neighboring Desmin-positive cells. Cells that were Desmin-positive but lacked connected Desmin-positive neighbors were considered isolated.

To evaluate statistical significance of the spatial distribution of Desmin-positive cardiomyocytes, null simulations were performed under two conditions. In the random-labeling (RL) model, cell positions were fixed, but Desmin-positive cells were randomly reassigned. In the complete spatial randomness (CSR) model, Desmin-positive and Desmin-negative cells were randomly distributed across the entire available area of the image while maintaining the observed total numbers of Desmin-positive and Desmin-negative cells. In both cases, simulations were repeated 999 times, and the distribution of simulated cells was compared to the observed cells in terms of the percentage of Desmin-positive cells that were isolated and the percentage of Desmin-positive cells that were found in clusters of ≥3 cells. We also directly assessed the *p* value in comparing the clustering observed to the clustering measured from the 999 simulations **(Fig. S8 A-C).**

To address the overall morphology of the clusters of Desmin-positive cardiomyocytes (**Fig. S8 D**), we randomly selected 25 images that each occupied less than 20% of the available ROI area (to allow unambiguous interpretation of cluster morphology). We then assessed eccentricity and circularity of the 25 clusters of cells (treating each cluster as a single object).

### 1.5 Device Fabrication:

The 3D-printed mold and detailed geometry of the post design used in this study are shown in **(Fig. S2 A, B)**. Over time, cells actively remodeled the surrounding collagen matrix, leading to progressive gel compaction. This was reflected by a marked reduction in tissue area (normalized to the micro-well's area) until day 7 (**Fig. S2 C, D**). Following formation, μHT performed contractile work against the elastic resistance of the posts, mimicking the mechanical strain experienced by ventricular tissue *in vivo*. μHTs exhibited synchronous, spontaneous contractions within the first week of culture, as indicated by post deflection (**Supplement video 1)**. Variability in tissue formation efficiency and size is a common limitation in micro-scale engineered heart tissues (2). In our original platform, the posts were 1 mm in height with a cylindrical shaft with 200 μm base diameter, and a flat cap with 300 μm diameter. In practice, tissues did not consistently climb up to the cap of the posts, which led to variability in tissue geometry and limited throughput (**Fig. S3 A)**. To promote uniform tissue formation and size across independent replicates, we designed adapted alternative "sloped" micropillar geometry that encouraged uniform "climbing" of the μHT to the top of the PDMS post, which was capped with an overhanging cap to prevent tissue detachment (**Fig. S3 B**). While this slope did increase post bending stiffness **(Fig. S3 A, B)**, it aided both tissue formation as well as device fabrication.

**1.6 FEM Simulation:**

Finite element modeling was performed in Ansys Workbench 2025 R1 (Static Structural module) to estimate post deflection under applied contractile forces. Material properties of PDMS were first defined in the Engineering Data of Workbench, with Young's modulus ranging from 640–3000 kPa and a Poisson's ratio of 0.495. Geometries of individual posts were designed in Autodesk Fusion 360 and imported into Workbench's design modeler as external files (.f3d). Models were solved using the Mechanical APDL solver, with PDMS properties assigned to the post geometry. A patch-conforming mesh with quadratic tetrahedral elements (10 μm size, SOLID187) was generated using the quad dominant method, resulting in 90,671 nodes and 52,659 elements for the "new design" geometry. Boundary conditions were defined by fixing the base of the post, while contractile force was applied along the positive x-axis on the external face of the post (to the right), within a region 100–300 μm below the inferior edge of the cap. To evaluate post deformation under different material properties, Young's modulus (0.64, 1.1, and 3.0 MPa), Poisson's ratio (0.40, 0.4495, and 0.499), and applied force were set as input parameters in a response surface optimization analysis. The maximum total deformation was defined as the output parameter. This design of experiments generated 15 design points, allowing to estimate deflection across the specified parameter space **(Fig. S3 A, B)**.

### 1.7 Exclusion Criteria:

All μHTs passed initial quality control based on cardiac troponin T (cTnT) positivity, with flow cytometry confirming >95% $cTnT^+$ cardiomyocyte purity for both isogenic control and CRYAB-R120G mutant lines (**Fig. S11 A**). This ensured that observed functional differences were not attributable to variations in differentiation efficiency. The second quality control involved verifying that tissues maintained proper morphology and climbed up to the cap of the posts, which is essential for reliable point-force contractility measurements. Tissues that did not meet this criterion were excluded from analysis (**Fig. S11 B, C**). Altogether, these quality control metrices led to a more consistent and narrower range of μHT physiologic behaviors.

### Supplementary Figures:

Figure S1

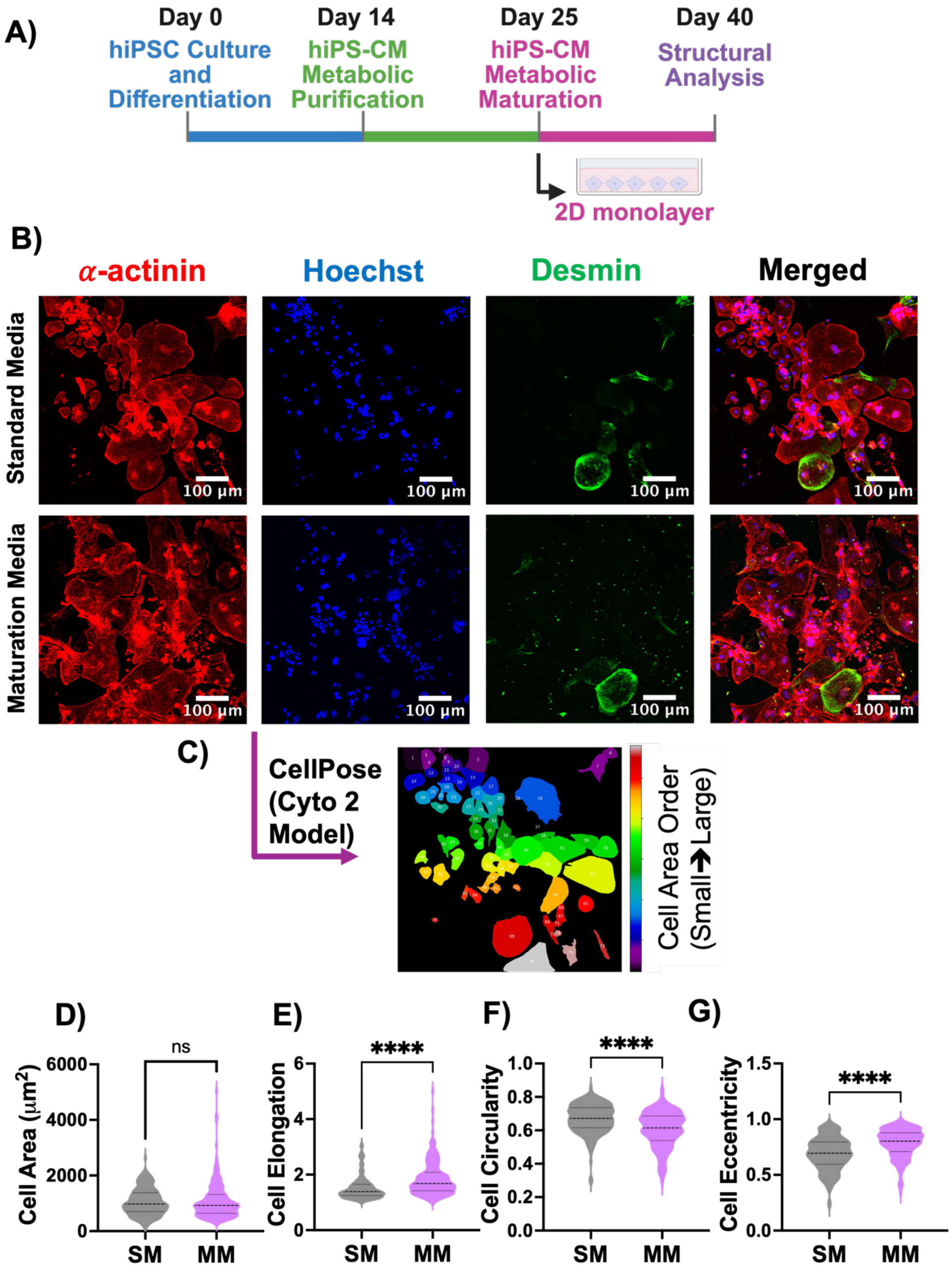


**Figure S1. Metabolic maturation alters hiPS-CM morphology without significantly changing Desmin expression. (A)** Schematic of the experimental timeline for generating hiPSC-derived cardiomyocytes (hiPS-CMs) in 2D monolayers. Cells were differentiated from hiPSCs (D0–D14), purified by metabolic selection (D14–D25), and cultured in either standard media (SM) or maturation media (MM) from D25–D40 before analysis. **(B)** Immunofluorescence staining for Desmin (green), α-actinin (red), and Hoechst (blue) in SM and MM cultured hiPS-CMs at D40. Scale bars: 100 μm. **(C)** Representative CellPose segmentation (Cyto2 model) of α-actinin/Hoechst-stained images with cells color-coded by area (small → large). Quantification of cell **(D)** area, **(E)** elongation, **(F)** circularity, and **(G)** eccentricity from α-actinin/Hoechst-stained images. MM-cultured hiPS-CMs exhibit increased eccentricity and elongation, decreased circularity with no significant difference in cell area compared to SM cultured cells. For all quantifications, data are presented as mean ± *SD*; n values represent individual cells pooled from 3 independent differentiations. *P* values by unpaired t-test, non-parametric, Mann-Whitney test; **** $p < 0.0001$, ns = not significant.

**Figure S2**

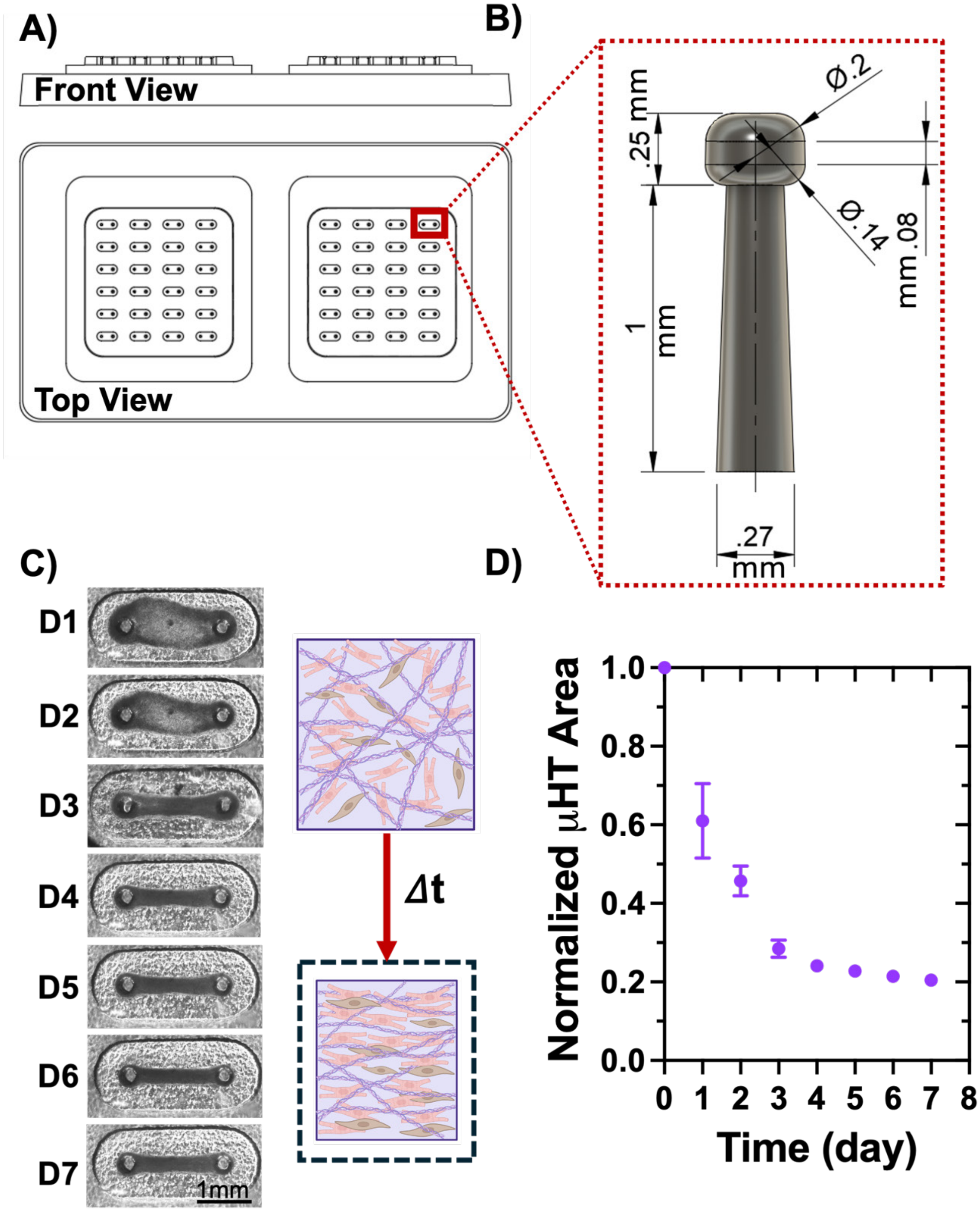


**Figure S2. Device design for 3D μHT formation and time-dependent tissue compaction. (A)** Schematic of the 3D μHT device showing front and top views of the multi-well format and **(B)** detailed dimensions of an individual PDMS post used for tissue anchoring. Posts are 1 mm tall with a 0.25 mm cap height, designed to support micro-tissue formation between paired posts.

**(C)** Representative brightfield images of μHTs from day 1 (D1) to day 7 (D7) after seeding, showing progressive tissue compaction. Insets illustrate fiber compaction and alignment over time (Δt). **(D)** Quantification of normalized μHT area shows a rapid decrease within the first 3 days, followed by stabilization by day 5–7. Data represent mean ± *SD* of tissues from 2 differentiation batches. Scale bar: 1 mm.

Figure S3

A)

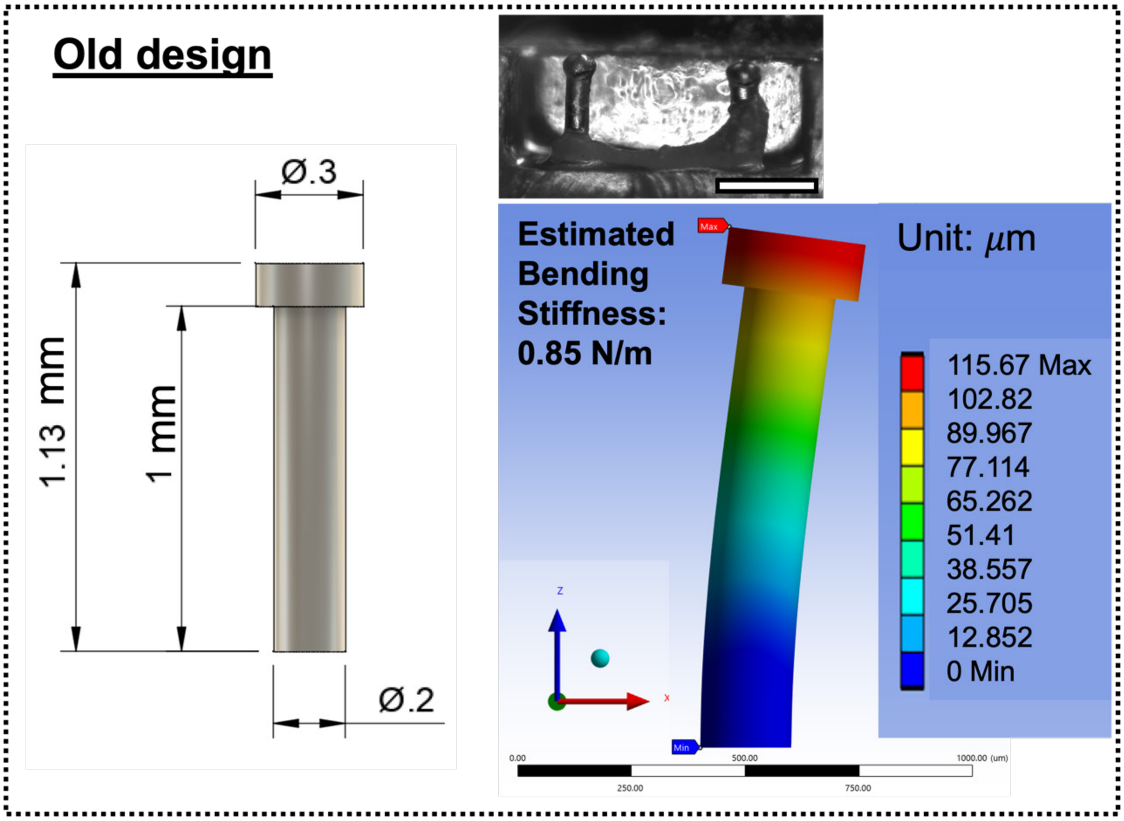
Old design
Ø.3
1.13 mm
1 mm
Ø.2
Estimated Bending Stiffness: 0.85 N/m
Unit: μm
115.67 Max
102.82
89.967
77.114
65.262
51.41
38.557
25.705
12.852
0 Min

B)

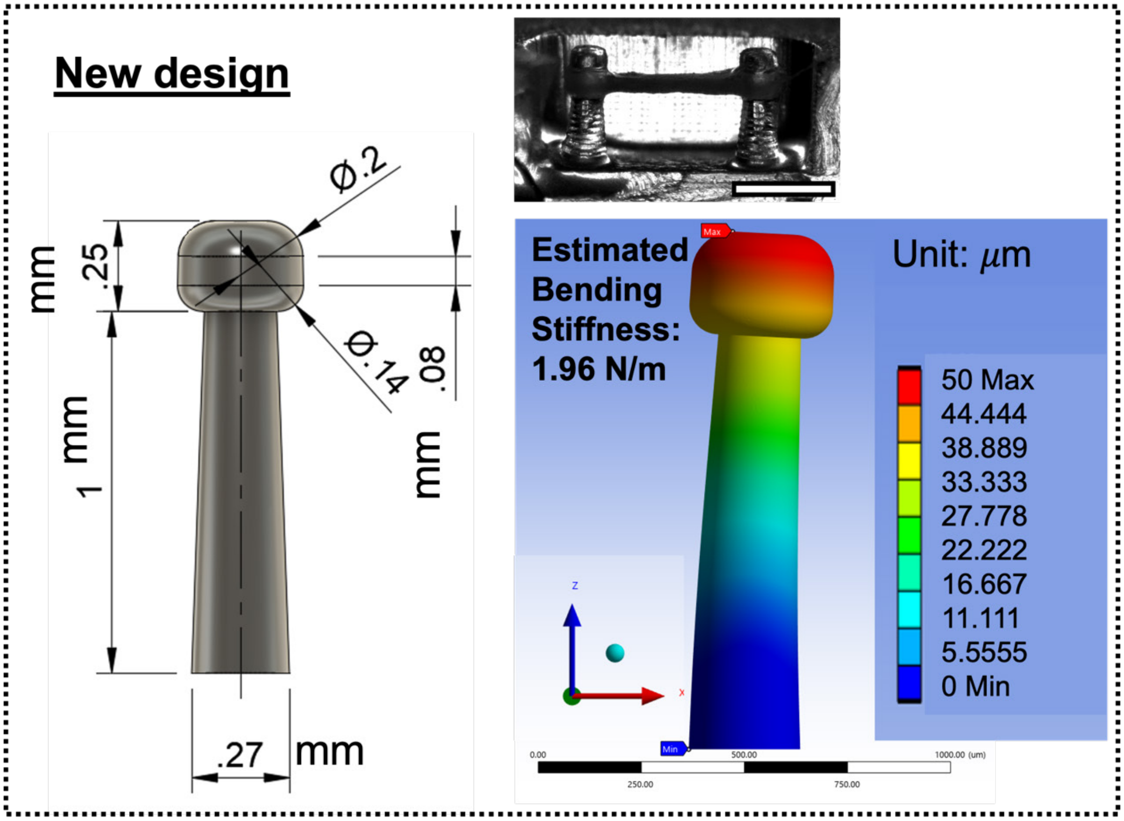
New design
Ø.2
.25 mm
1 mm
Ø.14
.08 mm
.27 mm
Estimated Bending Stiffness: 1.96 N/m
Unit: μm
50 Max
44.444
38.889
33.333
27.778
22.222
16.667
11.111
5.5555
0 Min

**Figure S3. Comparison of old and new PDMS post designs for consistent 3D µHT formation. (A)** Old post design: schematic showing dimensions (1.13 mm total height, 0.3 mm cap diameter, 0.2 mm shaft diameter) and representative brightfield image of front view µHT formed between posts. Finite element model (FEM) for the old design within stiff post condition (E = 3000kPa) is shown. **(B)** New post design: schematic showing dimensions (1 mm in height, with a 270 µm base diameter and a –2° taper) and representative brightfield image of µHT formed between posts. FEM simulation for the stiff post condition (E = 3000 kPa) is shown. The new design supports more robust tissue attachment and improved µHT formation compared to the old design. Scale bars: 1mm (brightfield images).

**Figure S4**

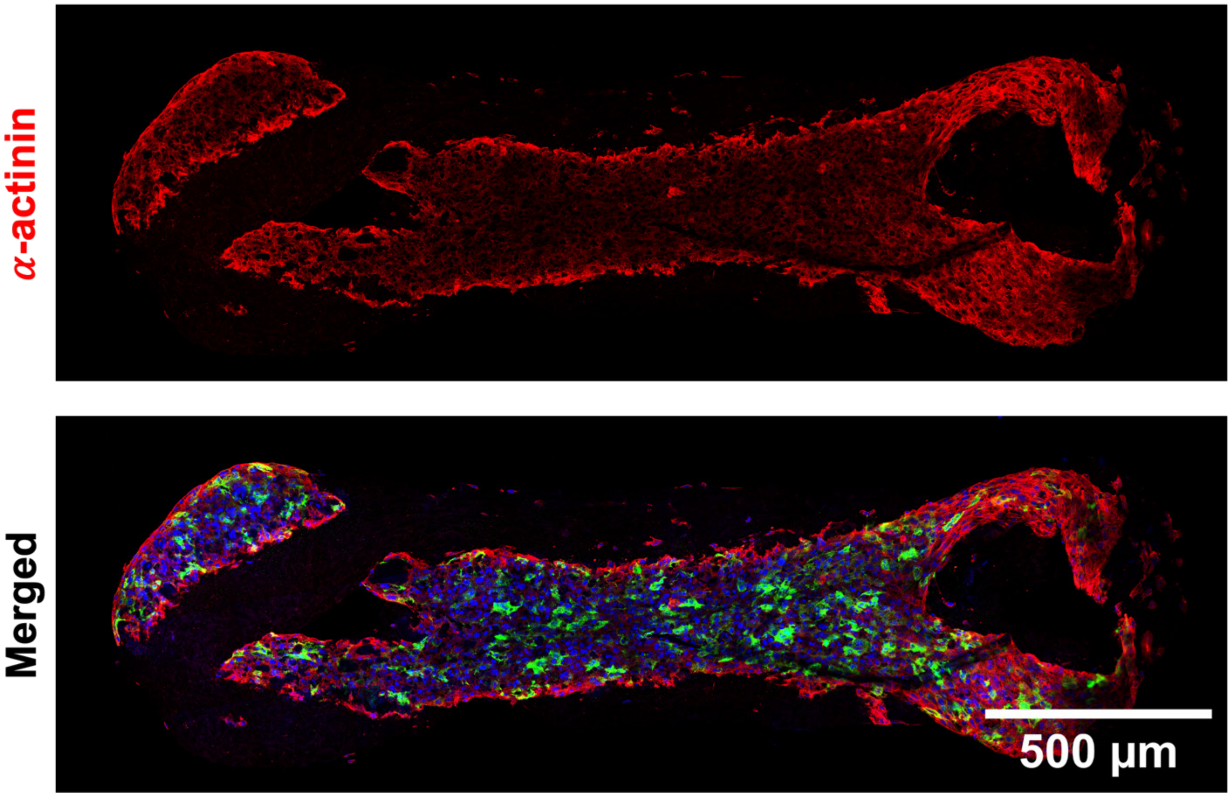


**Figure S4. Desmin is expressed throughout the μHT irrespective of local mechanical stress.** Immunofluorescence whole-mount staining for α-actinin (red), Desmin (green), and Hoechst (blue) of a 3D μHT. Desmin is uniformly distributed throughout the μHT independent of spatial proximity to mechanical anchor points. Scale bar: 500 μm.

Figure S5

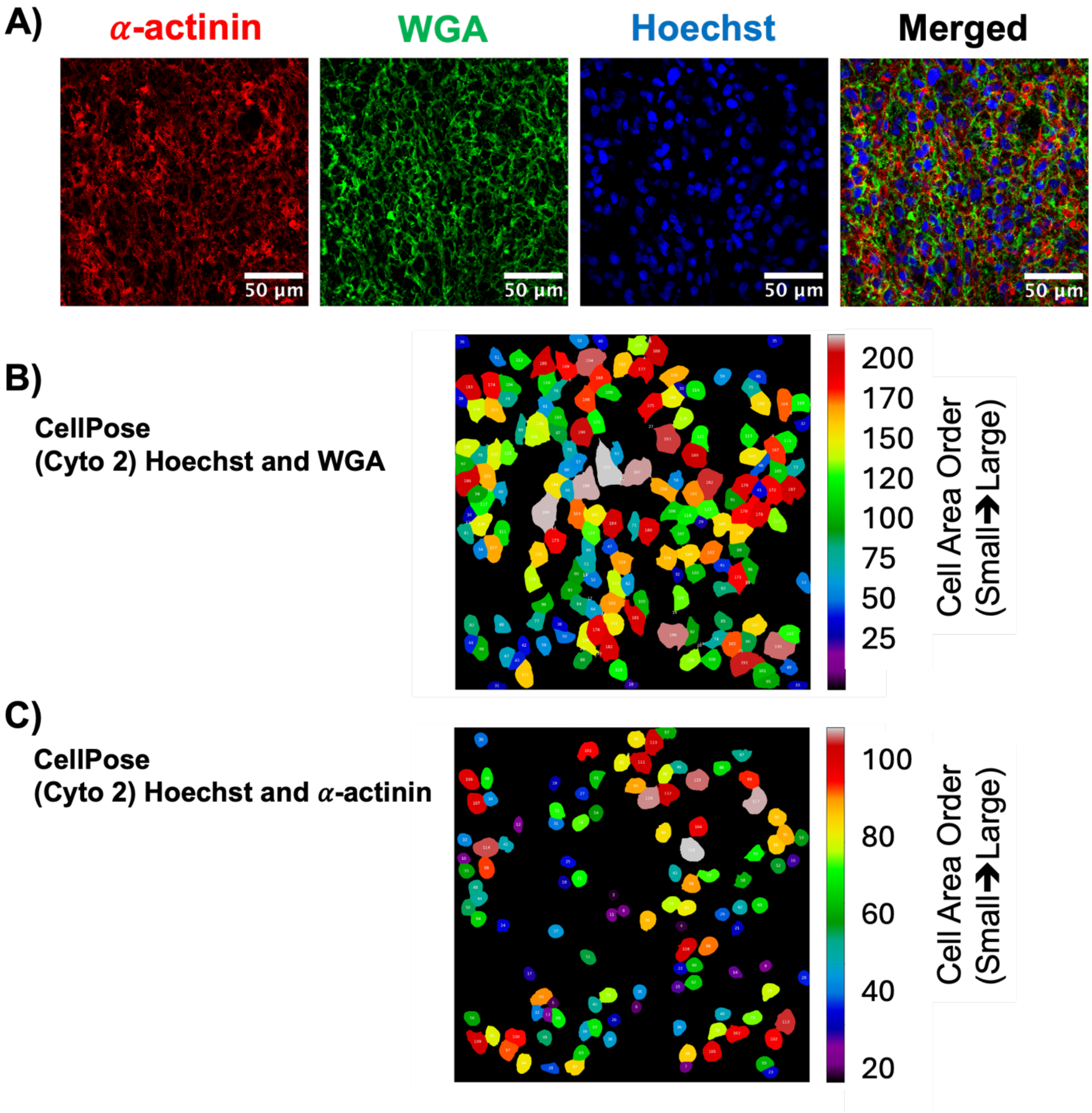


**Figure S5. Whole-mount 3D μHT imaging and segmentation analysis using CellPose (Cyto2 model). (A)** Representative immunofluorescence staining for α-actinin (red), WGA (green), and Hoechst (blue), with merged image shown. Scale bars: 50 μm. **(B)** Cell segmentation performed using the CellPose Cyto2 model with WGA and Hoechst images, showing identified cell boundaries color-coded from small to large cell area. **(C)** Cell segmentation performed using the

CellPose Cyto2 model with α-actinin and Hoechst images, showing identified cell boundaries color-coded from small to large cell area.

### Figure S6

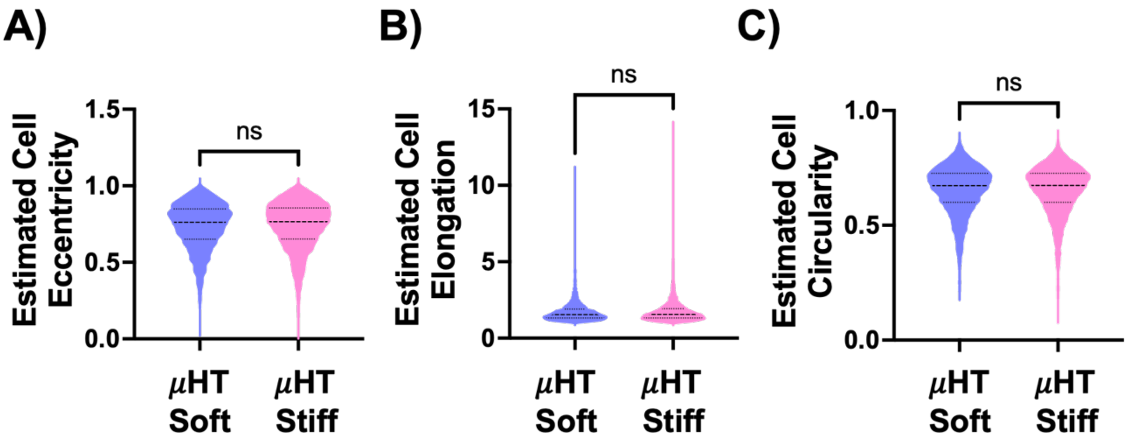


**Figure S6. 3D µHT culture between PDMS posts with increasing stiffness modulates hiPS-CM morphology in isogenic controls.** Quantification of estimated cell **(A)** eccentricity, **(B)** elongation, and **(C)** circularity from nuclei-stained µHTs, showing no significant difference in cellular morphology between stiff and soft posts. Data are presented as mean ± *SD*; n values represent individual cells pooled from 3 independent differentiation batches. *P* values by unpaired Student t-test, non-parametric, Mann-Whitney test; ns = not significant.

**Figure S7**

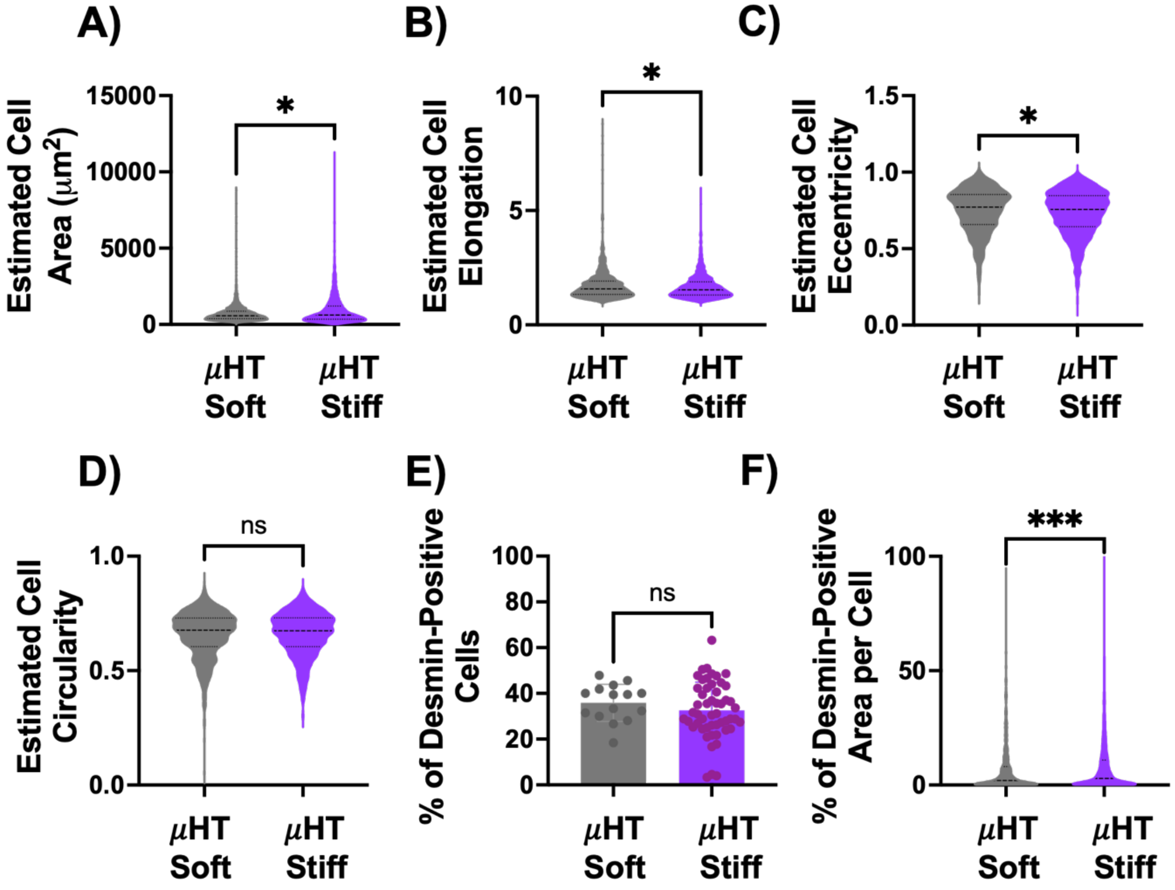


**Figure S7. 3D μHT culture between PDMS posts with increasing stiffness modulates morphology and Desmin organization in CRYAB-R120G mutant hiPS-CMs. (A)** Quantification of estimated cell area from nuclei-stained μHTs, showing significantly increased cell area on stiff posts compared to soft posts. **(B)** Quantification of estimated cell elongation, revealing decreased elongation on stiff posts. **(C)** Quantification of estimated cell eccentricity, showing significantly lower eccentricity on stiff posts. **(D)** Quantification of estimated cell circularity, showing no significant difference between stiff and soft posts. Data are presented as mean ± *SD*; n values represent individual cells pooled from 3 independent differentiation batches. *P* values were determined by unpaired Student's t-test, non-parametric, Mann–Whitney test; * $p < 0.05$, ns = not significant. **(E)** Quantification of the percentage of Desmin-positive cells, shows no significant difference between stiff and soft posts. Data are presented as mean ± *SD*; n values represent individual µHTs pooled from 3 independent differentiation batches. *P* values were determined by unpaired Student's t-test, parametric; ns = not significant. **(F)** Quantification of the percentage of Desmin-positive area per cell, showing a significant increase on µHTs within stiff posts compared to soft posts. Data are presented as mean ± *SD*; n values represent individual cells pooled from 3 independent differentiation batches. *P* values were determined by unpaired Student's t-test, non-parametric, Mann–Whitney test; * $p < 0.05$.

### Figure S8

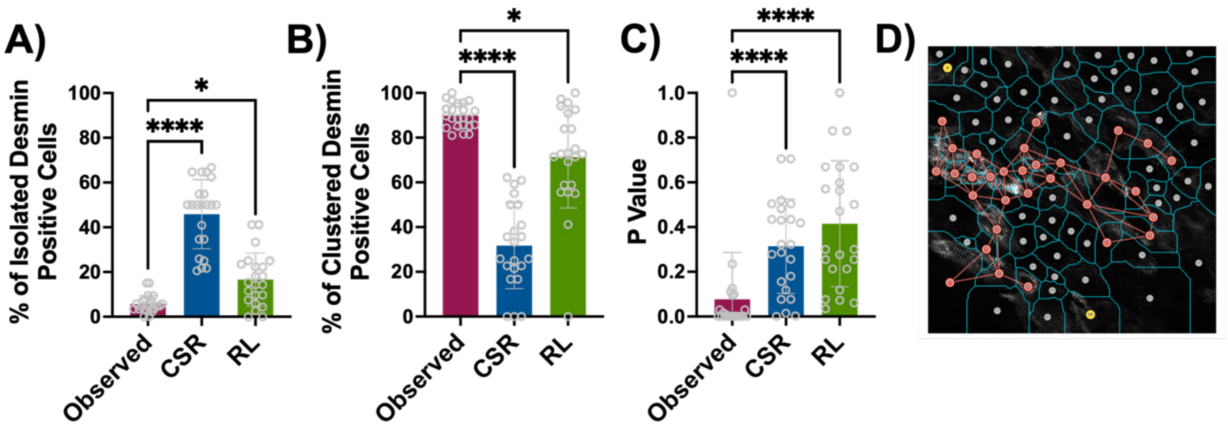


**Figure S8. Quantitative analysis of localized clustering of Desmin-positive hiPS-CM in µHT.** Quantification of the percentage of **(A)** isolated and **(B)** clustered Desmin-positive cells, and **(C)** corresponding *p*-values comparing observed distributions with null models of complete spatial randomness (CSR) and random labeling (RL). **(D)** Representative image of µHT segmentation with cell boundaries (blue) and line linking Desmin-positive cells (red) to its neighbors. Observed data demonstrated significant deviation from CSR, supporting non-random and clustered distribution of Desmin expression. Data are presented as mean ± *SD*; n values represent individual µHTs pooled from 3 independent differentiation batches. Statistical comparisons were performed by one-way ANOVA, non-parametric Kruskal–Wallis test; * $p < 0.05$, **** $p < 0.0001$.

**Figure S9**

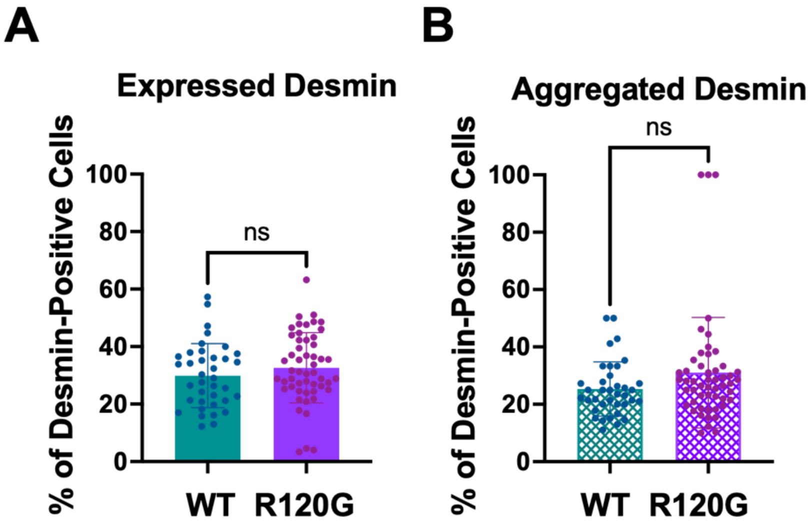


**Figure S9. Quantification of Desmin-positive cells and their aggregation across genotypes.** (**A)** Percentage of Desmin-positive cells (calculated as the number of Desmin-positive cells divided by the total number of cells). Data are presented as mean ± *SD*; n values represent individual µHT pooled from 3 independent differentiation batches. *P* values were determined by unpaired Student's t-test, parametric; ns = not significant. **(B)** Percentage of Desmin-positive cells that are aggregated (calculated as the number of Desmin-positive cells in aggregated form divided by Desmin-positive cells). Data are presented as mean ± *SD*; n values represent individual µHT pooled from 3 independent differentiation batches. *P* values were determined by unpaired Student's t-test, non-parametric, Mann–Whitney test; ns = not significant.

**Figure S10**

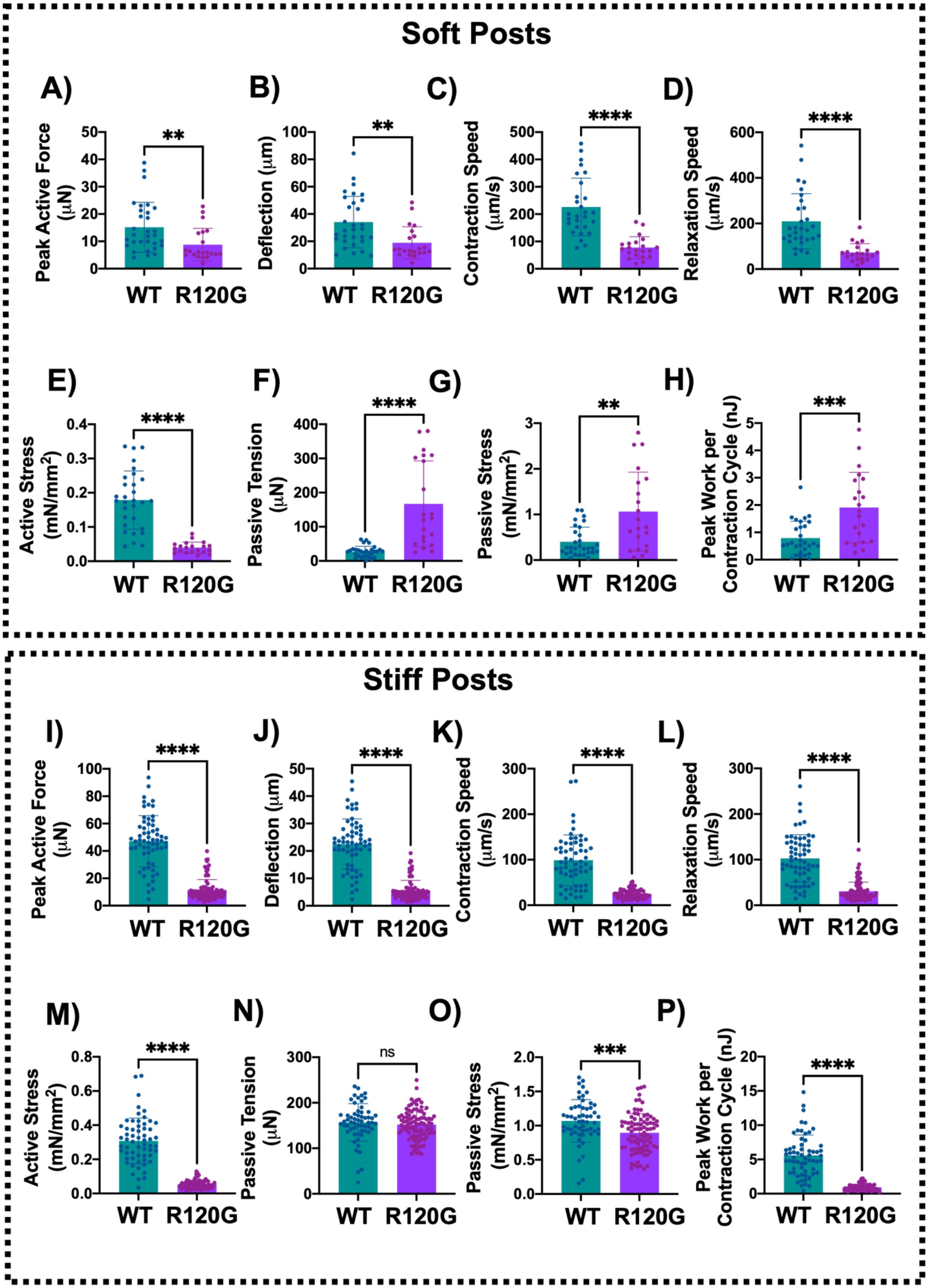

Soft Posts
A)
Peak Active Force (μN)
**
WT
R120G
B)
Deflection (μm)
**
WT
R120G
C)
Contraction Speed (μm/s)
****
WT
R120G
D)
Relaxation Speed (μm/s)
****
WT
R120G
E)
Active Stress (mN/mm$^2$)
****
WT
R120G
F)
Passive Tension (μN)
****
WT
R120G
G)
Passive Stress (mN/mm$^2$)
**
WT
R120G
H)
Peak Work per Contraction Cycle (nJ)
***
WT
R120G
Stiff Posts
I)
Peak Active Force (μN)
****
WT
R120G
J)
Deflection (μm)
****
WT
R120G
K)
Contraction Speed (μm/s)
****
WT
R120G
L)
Relaxation Speed (μm/s)
****
WT
R120G
M)
Active Stress (mN/mm$^2$)
****
WT
R120G
N)
Passive Tension (μN)
ns
WT
R120G
O)
Passive Stress (mN/mm$^2$)
***
WT
R120G
P)
Peak Work per Contraction Cycle (nJ)
****
WT
R120G

**Figure S10. Functional profiling of µHTs within soft and stiff PDMS posts reveals contractile deficit for CRYAB-R120G µHTs.** Quantification of individual contractile parameters from isogenic controls and CRYAB-R120G µHTs cultured on **(A–H) soft posts** (top panels) and **(I-P) stiff posts** (bottom panels). Parameters measured include: **(A, I)** peak active force (µN), **(B, J)** post deflection (µm), **(C, K)** contraction speed (µm/s), **(D, L)** relaxation speed (µm/s), **(E, M)** active stress ($mN/mm^2$), **(F, N)** passive tension (µN), **(G, O)** passive stress ($mN/mm^2$), and **(H, P)** peak work per contraction cycle (nJ). Data are presented as mean ± *SD*; n values represent individual µHTs pooled from 3 independent differentiation batches. *P* values were determined by unpaired Student's t-test, non-parametric Mann–Whitney test; ** $p < 0.01$, *** $p < 0.001$, **** $p < 0.0001$, ns = not significant.

**Figure S11**

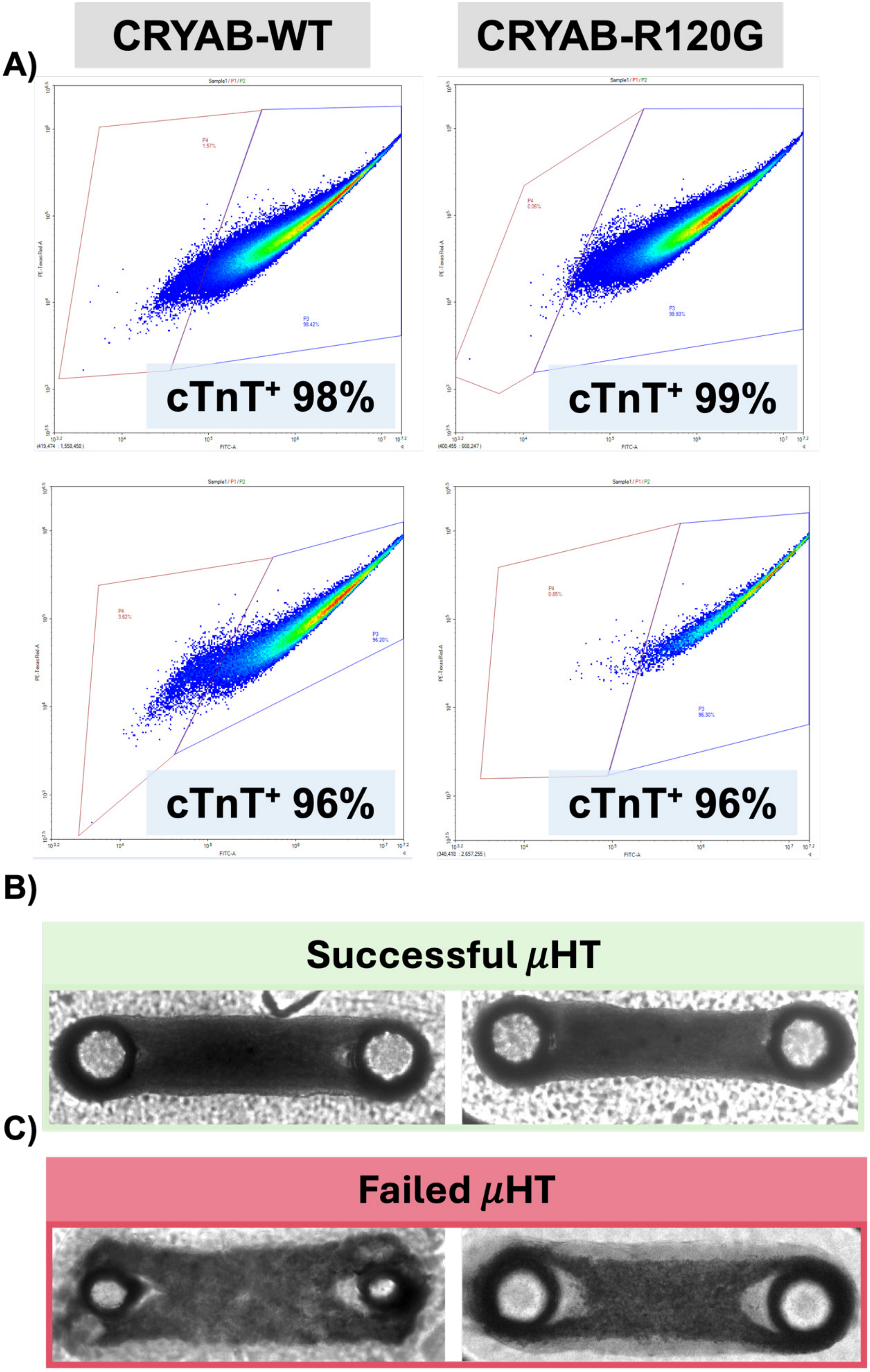

A)
CRYAB-WT
CRYAB-R120G
cTnT+ 98%
cTnT+ 99%
cTnT+ 96%
cTnT+ 96%
B)
Successful μHT
C)
Failed μHT

**Figure S11**. **Exclusion criteria for µHT experiments.** **(A)** Flow cytometry plots for cardiac troponin T (cTnT) expression in CRYAB-WT and CRYAB-R120G hiPS-CMs prior to µHT formation. Only preparations with >95% $cTnT^+$ cells were included in experiments (representative examples: 96–99% $cTnT^+$). **(B)** Representative brightfield images of µHTs passing inclusion criteria: tissues are compact, well-aligned, and climb up the PDMS posts ("successful µHT"). **(C)** Representative brightfield images of µHTs failing inclusion criteria: tissues show poor compaction, detachment, or failure to engage the PDMS posts ("failed µHT").

### Figure S12

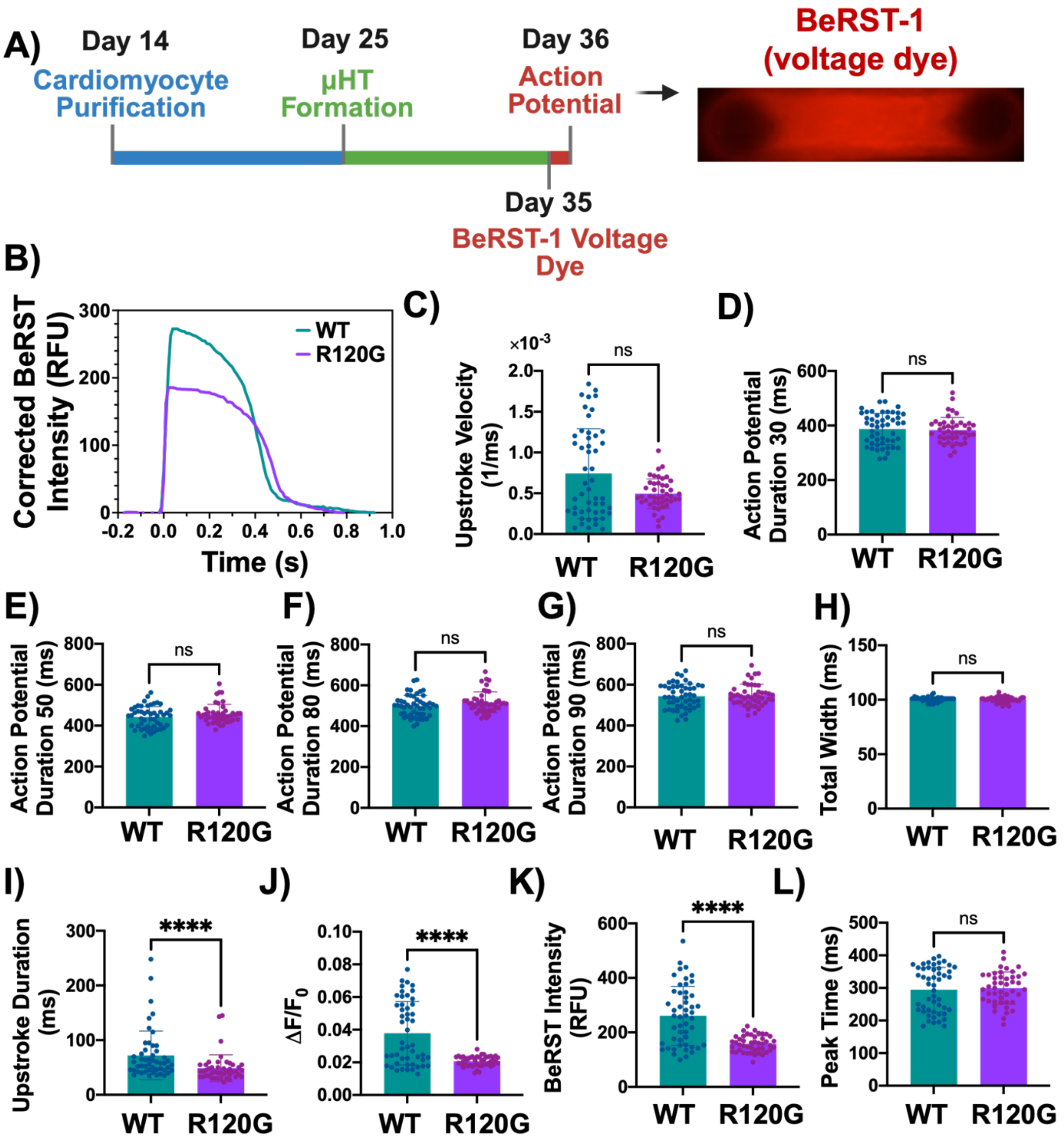


**Figure S12. Action potential properties of isogenic controls and CRYAB-R120G-μHTs measured by BeRST voltage-sensitive dye. (A)** Experimental timeline and representative image of μHT loaded with BeRST-1 (voltage dye). **(B)** Representative corrected BeRST-1 intensity traces from CRYAB-WT and CRYAB-R120G μHTs. Quantification of **(C)** upstroke velocity measured by normalized background-corrected BeRST-1 signal intensity ($\Delta F/F_0$) over upstroke duration time. Action potential duration at **(D)** 30% ($APD_{30}$), **(E)** 50% ($APD_{50}$), **(F)** 80% ($APD_{80}$), and **(G)** 90% ($APD_{90}$) repolarization were measured. Quantification of **(H)** total width (ms), **(I)** upstroke duration (ms), **(J)** normalized amplitude ($\Delta F/F_0$), **(K)** BeRST fluorescence intensity (RFU), and **(L)** peak time (ms). CRYAB-R120G µHTs show significantly reduced action potential amplitude with no significant change in action potential durations and upstroke velocity. Data are presented as mean ± *SD*; n values represent individual µHTs pooled from 3 independent differentiation batches. Statistical comparisons were performed by unpaired t-test for parametric data ($APD_{90}$) and Mann–Whitney test for non-parametric data; **** $p < 0.0001$, ns = not significant.

**Figure S13**

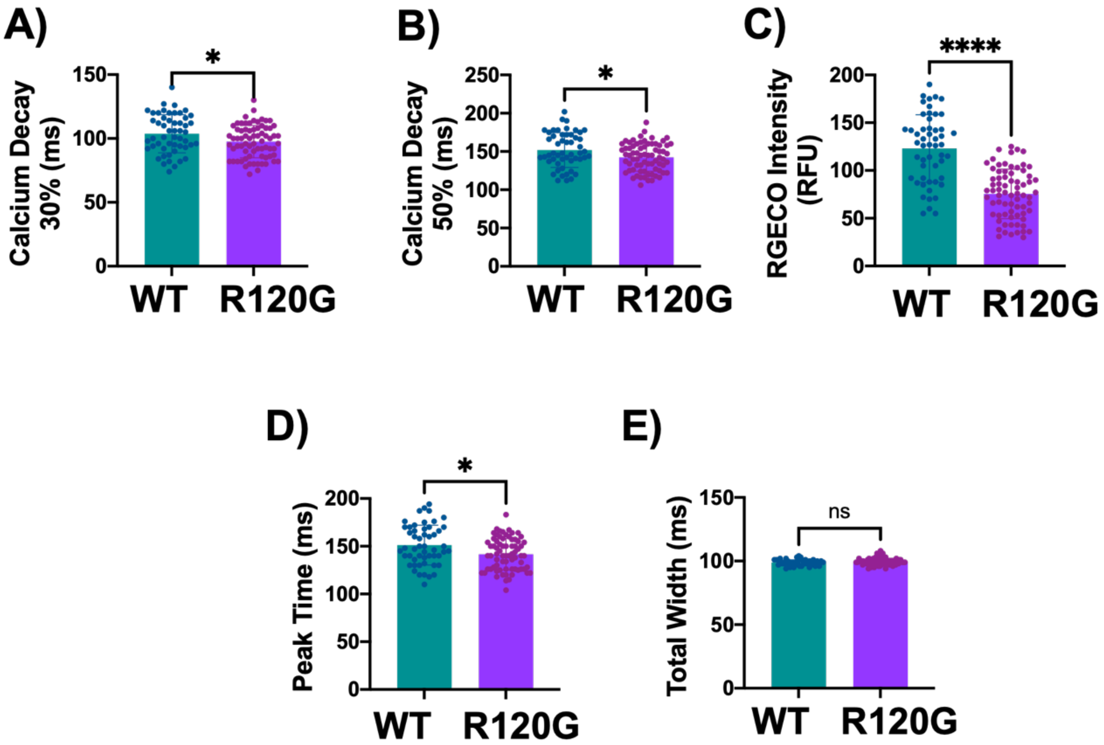


**Figure S13. Calcium handling defects in CRYAB-R120G µHTs revealed by RGECO1.2 imaging. (A–E)** Quantification of calcium transient parameters from isogenic controls and CRYAB-R120G µHTs expressing the genetically encoded calcium indicator RGECO1.2.

Quantification of **(A)** calcium decay 30% (ms), **(B)** calcium decay 50% (ms), **(C)** RGECO fluorescence intensity (RFU), **(D)** peak time (ms), and **(E)** total width (ms). Data are presented as mean ± *SD*; n values represent individual μHTs pooled from 3 independent differentiation batches. Statistical comparisons were performed by unpaired Student's t-test for parametric data (calcium decay 30% and calcium decay 50%) and by Mann–Whitney test for non-parametric data (RGECO intensity, peak time, and total width); * $p < 0.05$, **** $p < 0.0001$, ns = not significant.

**Figure S14**

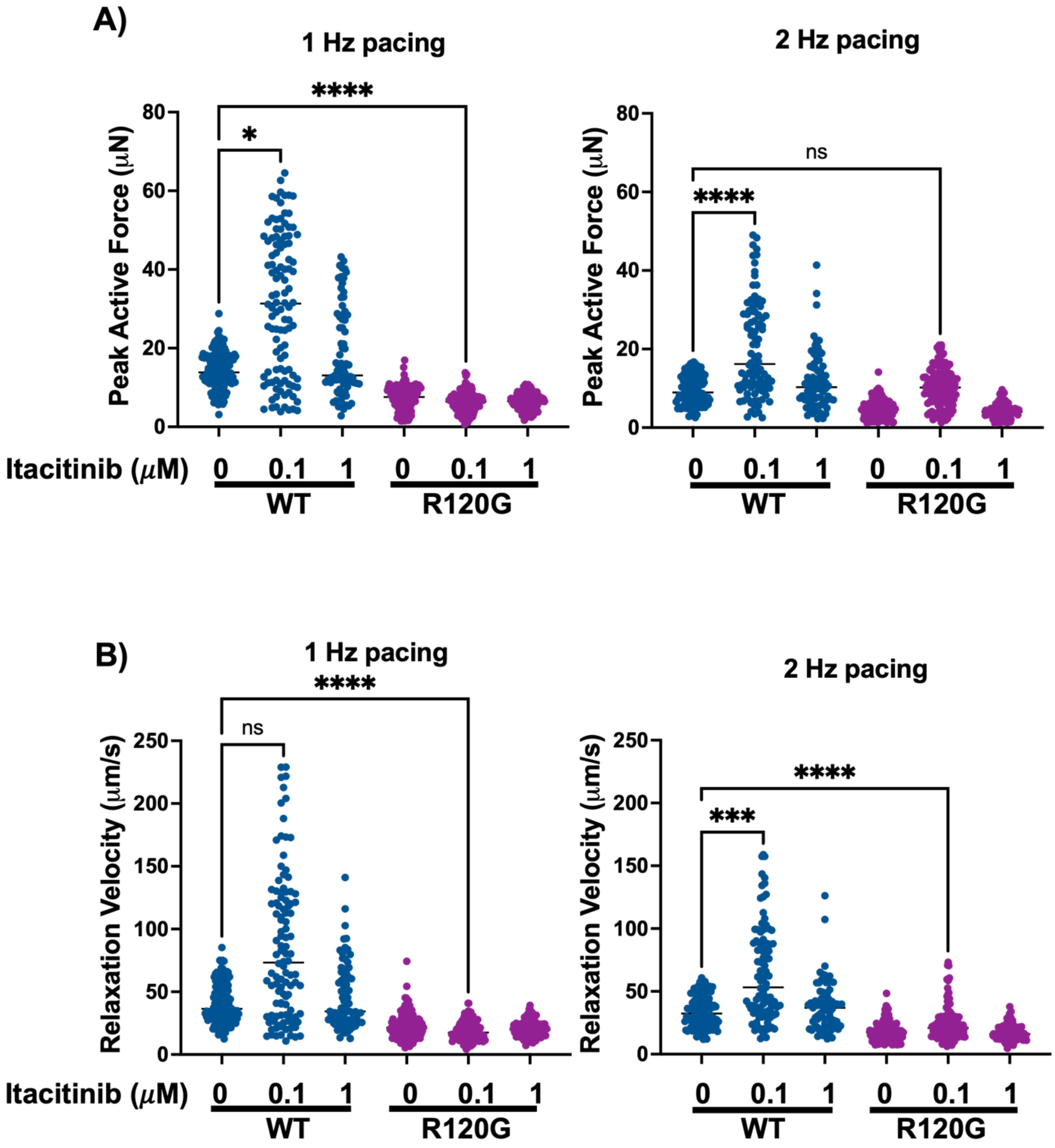


**Figure S14. Itacitinib treatment partially rescues contractile performance of CRYAB-R120G μHTs. (A)** Quantification of peak active force at 1 Hz (left) and 2 Hz (right) pacing. CRYAB-WT μHTs showed significantly increased absolute peak active force with 0.1 μM Itacitinib at 1 and 2 Hz. CRYAB-R120G μHTs showed consistently reduced absolute peak active force compared to

WT. However, treatment with 0.1 μM Itacitinib improved peak active force in CRYAB-R120G μHTs relative to its vehicle control, and levels did recover to those of WT vehicle controls at 2 Hz pacing (not 1 Hz pacing) but did not reach the isogenic controls after treating with 0.1 μM Itacitinib. **(B)** Quantification of relaxation velocity at 1 Hz (left) and 2 Hz (right) pacing. CRYAB-WT μHTs exhibited a significant increase in relaxation velocity at 0.1 μM Itacitinib compared to vehicle at both 1 and 2 Hz pacing, while R120G μHTs showed no improvement across conditions. Treatment with 0.1 μM Itacitinib improved relaxation velocity in CRYAB-R120G μHTs relative to its vehicle control, but levels did not recover to those of WT vehicle controls at both 1 and 2 Hz pacing. Data are presented as mean ± *SD*; n values represent individual μHTs pooled from ≥2 independent differentiations. *P* values determined by one-way ANOVA; non-parametric with Dunn's correction; **** $p < 0.0001$, *** $p < 0.001$, * $p < 0.05$, ns = not significant.

**Figure S15**

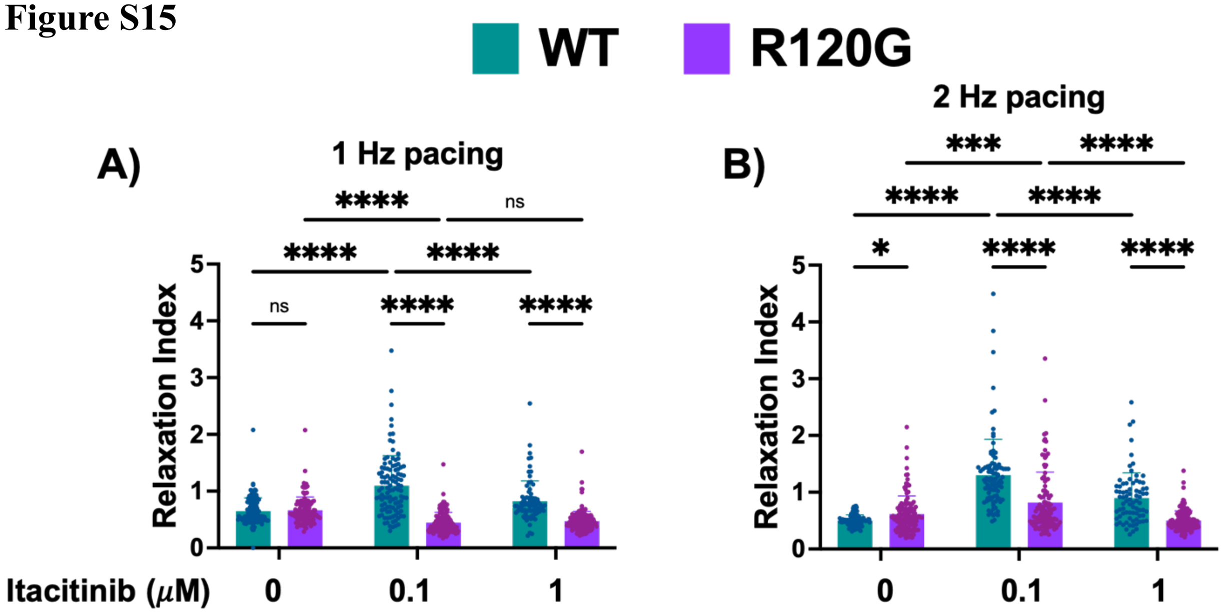

**Figure S15. Itacitinib partially rescued relaxation deficit in CRYAB-R120G µHTs.** Relaxation Index (ratio of relaxation velocity post-/pre-treatment) of CRYAB-WT and CRYAB-R120G µHTs treated with vehicle control, 0.1 µM, and 1 µM Itacitinib at **(A)** 1 Hz and **(B)** 2 Hz pacing frequency. 0.1 µM Itacitinib partially rescued relaxation velocity at 2 Hz in CRYAB-R120G µHTs; however, 1 µM Itacitinib worsened. Data are presented as mean ± *SD*; n values represent individual µHTs pooled from ≥2 independent differentiations. Data were analyzed by two-way ANOVA, followed by Holm–Šidák's multiple comparison test; **** $p < 0.0001$, *** $p < 0.001$, * $p < 0.05$, ns = not significant.
